# Isolation and characterisation of *Klebsiella* phages for phage therapy

**DOI:** 10.1101/2020.07.05.179689

**Authors:** Eleanor Townsend, Lucy Kelly, Lucy Gannon, George Muscatt, Rhys Dunstan, Slawomir Michniewski, Hari Sapkota, Saija J Kiljunen, Anna Kolsi, Mikael Skurnik, Trevor Lithgow, Andrew D. Millard, Eleanor Jameson

## Abstract

*Klebsiella* is a clinically important pathogen causing a variety of antimicrobial resistant infections in both community and nosocomial settings, particularly pneumonia, urinary tract infection and sepsis. Bacteriophage (phage) therapy is being considered as a primary option for the treatment of drugresistant infections of these types. We report the successful isolation and characterisation of 30 novel, genetically diverse *Klebsiella* phages. The isolated phages span six different phage families and nine genera, representing both lysogenic and lytic lifestyles. Individual *Klebsiella* phage isolates infected up to 11 of the 18 *Klebsiella* capsule types tested, and all 18 capsule-types were infected by at least one of the phages. Of the *Klebsiella*-infecting phages presented in this study, the lytic phages are most suitable for phage therapy, based on their broad host range, high virulence, short lysis period and given that they encode no known toxin or antimicrobial resistance genes. Importantly, when applied alone, none of the characterised phages were able to suppress the growth of *Klebsiella* for more than 12 hours, with some phages only able to suppress growth for 3 hours, likely due to inherent ease of *Klebsiella* to generate spontaneous phage-resistant mutants. This indicates that for successful phage therapy, a cocktail of multiple phages would be necessary to treat *Klebsiella* infections.

## Background

Bacteria of the *Klebsiella* genus are able to cause a variety of infections in both community and nosocomial settings including pneumonia, urinary tract infection (UTI), sepsis, wound infection, and infections in vulnerable populations including neonates and intensive care patients^1^. *Klebsiella* is also one of the most numerous secondary infection agents in COVID-19 patients, particularly those who have undergone ventilation^2–4^. Additionally, the sub-clinical carriage of *Klebsiella* is linked to cardiovascular disease^5, 6^ and inflammatory bowel disease^7^ *Klebsiella pneumoniae* is the most highly reported problematic pathogen from the genus; it has given rise to hypervirulent clones with extended virulence factors. Other *Klebsiella* species, including *K. oxytoca*^8^ and *K. variicola*^9^, are emerging pathogens, causing infections in immunocompromised patients. Whilst the arsenal of virulence factors make *Klebsiella* efficient pathogens, it is the high prevalence of antimicrobial resistance (AMR) mechanisms that complicates treatment and leads to high mortality^10^.

AMR represents a current threat to global health and security, fueled by our intensive use of antibiotics in both medicine and agriculture. *Klebsiella* is termed an ESKAPE pathogen on the World Health Organisation priority pathogens list^11^, gaining and transferring AMR genes, particularly in health-care settings^12^. The prevalence of resistance in *Klebsiella* has increased exponentially to most available antimicrobial drugs, and cases of pan-resistant *Klebsiella* are are now common around the world^13–16^. This poses a difficulty in treating these infections with existing antibiotics available in clinical settings^17^ and yet there are also major issues associated with vaccine development to prevent these infections^18^. The economic cost of *Klebsiella* outbreaks are high; in 2015 a Dutch hospital estimated the cost of an outbreak of multidrug-resistant (MDR) *K. pneumoniae* to be $804,263^19^. Alongside the economic consequences of these MDR *Klebsiella* infections, there is an increased risk of mortality^19, 20^. One potential alternative treatment to antibiotics are bacteriophages (phages)^21^.

Phages are natural killers of bacteria and phage therapy is emerging as a potential weapon against MDR bacterial infections^22, 23^. Phage therapy depends on preparedness, particularly in having a real or virtual “biobank” of phages suited for killing the bacteria responsible for common, AMR infections. The discovery of phages for therapeutic use is realistic given the low-costs of modern sequencing technologies and the classic technologies for characterization of phage virions^24^. Pre-therapeutic characterisation of phages is essential to mitigate side effects in patients, gain access to purified active phages in a timely manner, establish the best mode of delivery and the level of inactivation of phages by the immune system. It has been highlighted how awareness of broad aspects of phage biology can minimize potential side-effects during therapy^25^. Ultimately, these hurdles can be overcome with detailed understanding and characterisation of prospective phage genomes, structure and function.

In this paper we use genomic and imaging technologies to characterise novel phages isolated against clinical and environmental *Klebsiella* spp. Phages were discovered in a range of environmental scenarios including rivers, ponds, estuaries, canals, slurry, and sewage. The newly isolated phages were characterised using a combination of traditional and genomic approaches to understand their infection cycle, host range, and gene content. We present phages with siphovirus, myovirus, podovirus and inovirus morphologies, spanning six phage families and nine genera, of which the majority have lytic lifestyles. A number of these phages have potential use in phage therapy.

## Materials and Methods

### Bacterial Strains and Culture Conditions

The *Klebsiella* strains used in this work are listed in table 1. All culturing in liquid medium was performed with shaking (150 rpm) at 37 °C. All culturing was carried out in Lysogeny broth (LB), with the addition of 5 mM CaCl_2_ and 5 mM MgCl_2_ when culturing phages. The *Klebsiella* were originally isolated from clinical samples, except for two which were of environmental origin (table 1). The strains represented six species: *K. pneumoniae, K. oxytoca, K. quasipneumoniae, K. variicola, K. michagenesis* and *K. aerogenes*.

**Table 1.** Details of *Klebsiella* species and strains used in this study. Capsule types are given where applicable, alongside the origin of the strain and indication of use as an isolation host.

| Species | Strain | Capsule Type | Isolation Host? | Origin | Isolation source |
| --- | --- | --- | --- | --- | --- |
| *Klebsiella aerogenes* | 30053 | - | ✓ | DSMZ Culture Collection | sputum |
| *Klebsiella michiganensis* | 25444 | O1v1 | ✗ | DSMZ Culture Collection | toothbrush holder |
| *Klebsiella oxytoca* | 170748 | O1v1 | ✗ | Clinical Isolate | catheter specimen urine |
|  | 5175 | KL29 | ✗ | DSMZ Culture Collection | pharyngeal tonsil |
|  | 25736 | KL74 | ✓ | DSMZ Culture Collection | case of pneumonia |
|  | 170821 | OL104 | ✓ | Clinical Isolate | Urine |
|  | 171266 | OL104 | ✗ | Clinical Isolate | urostomy urine |
| *Klebsiella pneumoniae* | 170958 | KL28 | ✓ | Clinical Isolate | urine |
|  | 171304 | KL144 | ✗ | Clinical Isolate | catheter specimen urine |
|  | 13440 | KL38 | ✓ | NCTC Culture Collection | clinical |
|  | 13442 | KL110 | ✗ | NCTC Culture Collection | hospital, Italy |
|  | 30104 | KL3 | ✓ | DSMZ Culture Collection | human blood |
|  | 13465 | KL57 | ✗ | NCTC Culture Collection | clinical |
|  | 170820 | KL158 | ✗ | Clinical Isolate | urine |
|  | 16358 | KL4 | ✗ | DSMZ Culture Collection | human, nose |
|  | 170723 | KL2 | ✓ | Clinical Isolate | urine |
|  | 171167 | KL2 | ✗ | Clinical Isolate | urine |
|  | 13443 | KL2 | ✗ | NCTC Culture Collection | clinical |
|  | 13882 | KL64 | ✗ | ATCC Culture Collection | water |
|  | 13439 | KL14 | ✓ | NCTC Culture Collection | outbreak strain |
| *Klebsiella variicola* | W12 | KL14 | ✗ | Environmental Isolate | soil |
|  | 15968 | KL16 | ✓ | DSMZ Culture Collection | banana root |
| *Klebsiella quasipneumoniae* | 28211 | KL35 | ✓ | DSMZ Culture Collection | human blood |
|  | 700603 | KL53 | ✓ | ATCC Culture Collection | urine |

### Klebsiella *capsule typing*

Kaptive Web was used and determined that the 24 *Klebsiella* strains belonged to 18 different capsule types (table 1). Three *K. pneumoniae* strains were identified as capsule type KL2, while two *Klebsiella* sp. were each typed as O1v1, OL104 and KL14. All other capsule types were unique, *K. aerogenes* 30053 could not be typed.

### Phage Isolation

Phages were isolated from water samples from various sources, listed in table 2. Phages were named using the ICTV binomial system of viral nomenclature^26^. The water samples were filtered through 0.2 μm pore size syringe filters to remove debris and bacteria. Phages were then isolated by enrichment: 2.5 mL of filtered water sample was added to 2.5 ml nutrient broth, containing 5 mM CaCl_2_ and 5 mM MgCl_2_, and inoculated with 50 μL of overnight-grown *Klebsiella*. This enrichment culture was then incubated overnight at 37 °C, centrifuged, and the supernatant filtered through a 0.2 μm pore size filter to remove cells. This filtrate was serially diluted down to 10^-11^ in LB and used in an overlay agar plaque assay. Briefly, 50 μl of each serial dilution was mixed with 0.5 mL of a single *Klebsiella* strain in the logarithmic growth phase (OD_600_ 0.2) and incubated at room temperature for 5 min. To each serial dilution/cell mix, 2.5 mL of cooled, molten LB agar (0.4 % weight/volume) was added and mixed by swirling. The molten agar mix was poured onto 1 % LB agar plates. All overlay agar plates were allowed to set, then inverted and incubated overnight at 37 °C. From the plaque assay plates single plaques were identified, picked using a pipette tip, mixed with 50 μL of LB, and filtered through a 0.22 μm spin filter (Costar Spin-X, Corning, UK). The filtrate underwent two further rounds of plaque assay to ensure that phage isolates were the result of a single clonal phage.

**Table 2.** Phage isolate details. Lab ID refers to the laboratory identification number, source of isolation, indicates where the water sample was collected for phage enrichment and isolation and the strain of isolation indicates *Klebsiella* sp. strain on which 3 rounds of plaque assay isolation were performed. Accession numbers refers to associated project accession numbers assigned by the ENA for each phage.

| Phage Name | Lab ID | Source of isolation |  | Strain of isolation | Accession number |
| --- | --- | --- | --- | --- | --- |
| Klebsiella phage vB_KaS-Benoit | 1 | Estuary | Jelitkowo, Poland | *Klebsiella aerogenes* DSM 30053 | PRJEB39773 |
| Klebsiella phage vB_KaS-Veronica | 2 | Marine canal | Grand canal, Venice, Italy | *Klebsiella aerogenes* DSM 30053 | PRJEB40165 |
| Klebsiella phage vB_KvM-Eowyn | 4 | Estuary | Jelitkowo, Poland | *Klebsiella variicola* DSM 15968 | PRJEB40131 |
| Klebsiella phage vB_KaS-Gatomon | 6 | Marine canal | Grand canal, Venice, Italy | *Klebsiella aerogenes* DSM 30053 | PRJEB40159 |
| Klebsiella phage vB_KaS-Ahsoka | 7 | Slurry | Slurry tank, UK | *Klebsiella aerogenes* DSM 30053 | PRJEB40160 |
| Klebsiella phage vB_KppS-Samwise | 8 | Slurry | Slurry tank, UK | *Klebsiella pneumoniae* DSM 30104 | PRJEB40161 |
| Klebsiella phage vB_KppS-Totoro | 10 | Estuary | Jelitkowo, Poland | *Klebsiella pneumoniae* DSM 30104 | PRJEB40166 |
| Klebsiella phage vB_KoM-Pickle | 12 | Estuary | Jelitkowo, Poland | *Klebsiella oxytoca* DSM 25736 | PRJEB40176 |
| Klebsiella phage vB_KppS-Ponyo | 19 | River | Gneiwkowo, Poland | *Klebsiella pneumoniae* DSM 30104 | PRJEB40167 |
| Klebsiella phage vB_KppS-Jiji | 27 | Pond | Gneiwkowo, Poland | *Klebsiella pneumoniae* DSM 30104 | PRJEB40168 |
| Klebsiella phage vB_KppS-Raw | 33 | Sewage - raw | Spernal sewage works, UK | *Klebsiella pneumoniae* DSM 30104 | PRJEB40132 |
| Klebsiella phage vB_KppS-Storm | 34 | Sewage - storm tank | Spernal sewage works, UK | *Klebsiella pneumoniae* DSM 30104 | PRJEB40169 |
| Klebsiella phage vB_KppS-Ant | 35 | Sewage - anoxic sludge | Spernal sewage works, UK | *Klebsiella pneumoniae* DSM 30104 | PRJEB40148 |
| Klebsiella phage vB_KqM-LilBean | 36 | Sewage - raw | Spernal sewage works, UK | *Klebsiella quasipneumoniae* DSM 28211 | PRJEB40171 |
| Klebsiella phage vB_KqM-Bilbo | 38 | Sewage - raw | Spernal sewage works, UK | *Klebsiella quasipneumoniae* DSM 28211 | PRJEB40172 |
| Klebsiella phage vB_KqM-Westerburg | 39 | Sewage - raw | Spernal sewage works, UK | *Klebsiella quasipneumoniae* DSM 28211 | PRJEB40173 |
| Klebsiella phage vB_KpP-Yoda | 43 | Sewage - storm tank | Spernal sewage works, UK | *Klebsiella pneumoniae* 170723 | PRJEB40162 |
| Klebsiella phage vB_KqP-Goliath | 44 | Sewage - raw | Spernal sewage works, UK | *Klebsiella quasipneumoniae* DSM 700603 | PRJEB40163 |
| Klebsiella phage vB_KpP-Screen | 46 | Sewage - sieve | Spernal sewage works, UK | *Klebsiella pneumoniae* 170723 | PRJEB40164 |
| Klebsiella phage vB_KppS-Eggy | 49 | Sewage - anoxic sludge | Spernal sewage works, UK | *Klebsiella pneumoniae* DSM 30104 | PRJEB40146 |
| Klebsiella phage vB_KppS-Pokey | 50 | Sewage - anoxic sludge | Spernal sewage works, UK | *Klebsiella pneumoniae* DSM 30104 | PRJEB40147 |
| Klebsiella phage vB_KppS-Anoxic | 52 | Sewage - anoxic sludge | Spernal sewage works, UK | *Klebsiella pneumoniae* DSM 30104 | PRJEB40170 |
| Klebsiella phage vB_KoM-Liquor | 61 | Sewage - mixed liquor | Spernal sewage works, UK | *Klebsiella oxytoca* 170821 | PRJEB40174 |
| Klebsiella phage vB_KpM-Milk | 62 | Sewage - mixed liquor | Spernal sewage works, UK | *Klebsiella oxytoca* 170821 | PRJEB40175 |
| Klebsiella phage vB_KoM-Flushed | 63 | Sewage - mixed liquor | Spernal sewage works, UK | *Klebsiella pneumoniae* 170958 | PRJEB40181 |
| Klebsiella phage vB_KpM-Wobble | 64 | Sewage - mixed liquor | Spernal sewage works, UK | *Klebsiella pneumoniae* 170958 | PRJEB40182 |
| Klebsiella phage vB_KpM-Mild | 65 | Sewage - mixed liquor | Spernal sewage works, UK | *Klebsiella pneumoniae* DSM 13439 | PRJEB40177 |
| Klebsiella phage vB_KpM-KalD | 67 | Sewage - mixed liquor | Spernal sewage works, UK | *Klebsiella pneumoniae* DSM 13439 | PRJEB40178 |
| Klebsiella phage vB_KoM-MeTiny | 68 | Sewage - mixed liquor | Spernal sewage works, UK | *Klebsiella oxytoca* DSM 25736 | PRJEB40179 |
| Klebsiella phage vB_KpM-SoFaint | 70 | Sewage - mixed liquor | Spernal sewage works, UK | *Klebsiella pneumoniae* DSM 13440 | PRJEB40180 |

### DNA Extraction

DNA was extracted using the Phenol Chloroform Method^27^. Briefly, phage lysates were concentrated using a protein column with a 30 kDa cut-off. 750 μL of concentrated phage was treated with DNase I and Proteinase K, prior to phenol-chloroform, then overnight precipitation with ammonium acetate and ethanol at −20 °C. The DNA was resuspended in 50 μL of molecular grade water.

For bacteriophage which had a high background level of protein contamination from the host strain, Norgen Phage DNA Isolation Kit was used following the manufacturer’s instructions. To assess the quantity and quality of the isolated DNA for sequencing, both a spectrophotometer-based method and Qubit were used.

### Genome Sequencing

Sequencing was performed by MicrobesNG (Birmingham, UK), briefly; genomic DNA libraries were prepared using Nextera XT Library Prep Kit (Illumina, San Diego, USA) following the manufacturer’s protocol with modifications: 2 ng of DNA were used as the input, and a PCR elongation time of 1 min. DNA quantification and library preparation were carried out on a Hamilton Microlab STAR automated liquid handling system. Pooled libraries were quantified using the Kapa Biosystems Library Quantification Kit for Illumina, on a Roche light cycler 96 qPCR machine. Libraries were sequenced on the Illumina HiSeq using a 250 bp paired end protocol.

### Bioinformatics

Contig and genome assembly was carried out by MicrobesNG; reads were trimmed with Trimmomatic 0.30, sliding window quality cutoff of Q15^28^, SPAdes (v3.7) was used for *de novo* assembly^29^. Genomes were annotated using Prokka^30^, a custom database of phage genomes extracted from Genbank was used as previously described^31^. The capsule types of the *Klebsiella* strain genomes were predicted using Kaptive^32^.

To determine the taxonomy of our phages, the genomes were added to VIPtree^33^. Phage isolate genomes were then subject to the BLASTn and tBLASTn against NCBI. Average nucleotide identity (ANI) of the closest genomes identified were compared to our phages using orthoANI (https://www.ezbiocloud.net/tools/ani)^34^. Genomes with an ANI >95% were designated as the same species^35^. For genus-level clustering a shared protein network analysis was performed using vConTACT2 (v0.9.13)^36^ with all phage genomes available at the time (May 2020)^31^. The resulting network graph was visualised and annotated within Cytoscape (v3.8.0)^37^. Finally, multiple sequence alignments were performed using MAFFT (v7.271)^38^ on the DNA polymerase, large terminase subunit and major capsid proteins of each phage isolate genus with the most closely related phage proteins, identified. Phylogenetic trees were constructed with RaxML (v8.2.4)^39^ with 1000 bootstrap calculations using the GAMMA model of heterogeneity and the maximum-likelihood method based on the JTT substitution matrix. Subsequent trees were visualized and annotated in R (v3.6.1) using the ggtree (v1.16.6)^40, 41^ and phytools (v0.7-70) packages^42^.

Putative structural genes from the phage isolates were analysed for the presence of predicted depolymerase, enzymatic domains or features common to characterised depolymerase proteins. Each predicted gene product was analysed by BLASTP (v2.10.0; https://blast.ncbi.nlm.nih.gov/Blast.cgi), Pfam HMMER (v3.3; https://www.ebi.ac.uk/Tools/hmmer/) and HHpred (v33.1; https://toolkit.tuebingen.mpg.de/) using the default settings. Sequences from biochemically characterised depolymerase proteins that target *Klebsiella* spp. (table S1) or the putative depolymerases from our phage isolates (table S2) were used for analysis. Sequences were aligned with Muscle (v3.8.31)^43^ using SeaView (v4)^44^. Phylogenetic tree construction was performed with MegaX^45^ with 500 bootstrap calculations using the LG model. Tree topology searches were performed using a combination of NNI and NJ/BioNJ. The tree was subsequently visualised and annotated using iTOL(v4)^46^.

### Host Range Testing

Host range was tested by plating 5 μL of phage stock serial dilutions on to bacterial lawn, in 0.4 % overlay agar. Zones of clearing, indicating cell lysis were recorded: visible plaques, clearing at high phage titres, reduced lawn at high phage titres, turbid lawn with neat phage, or no effect.

### Plaque Formation and Morphology

Pure phage stocks were plated using the overlay agar plaque assay method, as described above, on their isolation host. Plates were incubated overnight at 37 °C to allow plaques to form. Plaque morphology was noted and photographs taken.

### Transmission Electron Microscopy

Pure phage stocks were imaged on formvar/carbon-coated copper grids (Agar Scientific Ltd, UK). To prepare grids for transmission electron microscope (TEM), the grids were glow-discharged for 1 min under vacuum. Following this, 5 μL of phage stock was applied to grid and incubated for 1.5 min at room temperature. The grid was then blotted to remove excess liquid. A drop of 2 % uranyl acetate was then applied to the grid and incubated for 1 min, before blotting off. The uranyl acetate staining was repeated four times. The grid was allowed to air dry after the final blot. Stained phage grids were imaged on a JEOL 2100Plus TEM. The morphology of the phage particles was visualised in ImageJ, phage particle capsids and tails were measured using the measure function. A median of thirty phage particles were used to calculate phage measurements.

### Lysis Period

*Klebsiella* cultures in the exponential growth phase were normalised to an OD_600nm_ of 0.2, using a spectrophotometer and phage lysates were diluted 1:4. The OD_600nm_ was measured every 5 min for 16 hr. Growth was compared to a positive control culture without the addition of phages. The lysis period was calculated by measuring the time from phage addition to a drop in culture OD_600nm_, relative to the positive control, indicating bacterial cell lysis.

### Virulence Index

The two virulence metrics the virulence Index (VP) and MV50 were calculated based on the protocol described by Storms and others^47^. Briefly, bacterial cultures were grown to exponential phase and then adjusted (as described for lysis period) to an optical density equivalent to 1 x 10^8^ cfu/mL. In a 96-well plate, phages were serially diluted from 1 x 10^8^ pfu/mL to 10 pfu/mL in 100 μL volumes. A bacterial culture was then added in equal volume (100 μL) to the phage dilution, resulting in multiplicity of infections (MOIs) from 1 to 10^-7^. The optical density of the 96-well plate was read at 600 nm at 5 min intervals for 18 hours. To calculate virulence indices, the area under the curve was calculated for both the bacterial only control and at each phage MOI, from time of initial infection until the exponential growth stage. The VP at each MOI was calculated following the method described^47^ using RStudio (version 1.1.463). The two virulence metrics capture different aspects of infection: VP is a quantified measure of the virulence of a phage against a bacterial host on a scale of 0-1 (from 0, no reduction in bacterial growth to 1, instantaneous complete killing); and the MV50 calculates the theoretical MOI at which a phage achieves a VP of 0.5 (half the theoretical maximum virulence).

### Data Visualisation

Resulting graphs were visualized in R (v3.6.1) implemented through RStudio (v1.1.456)^48^ using the ggplot2 (v3.3.2) package^49^, with a custom colour-blind colour palette generated from ColorBrewer (https://colorbrewer2.org).

## Results

### Sequence Similarity to Known Phages

The 30 *Klebsiella* phages were purified by multiple rounds of plating and genome sequencing showed a genome size range from 16,548 bp to 268,500 bp. The *Klebsiella* phage genomes represented nine diverse, distinct genera, as determined by VIPtree (figure 1) and vConTACT2 (figure 2). Genome similarities between our phage isolates and known phages were determined using a combination of VIPtree and BLAST (table 3). Phage isolates were grouped based on genus-level similarity into groups A-I, referred to by their genera or closest level of identifiable taxonomy: *Nonagvirus*, unclassified family/genus, *Tempevirinae* unclassified, *Myoviridae* unclassified, *Drulisvirus*, *Sugarlandvirus, Taipeivirus, Slopekvirus* and *Jiaodavirus*. The phage isolates of groups B, C and D displayed lower sequence similarities to previously identified phages, hence they were not classified into known genera. The sequence data was deposited in the ENA, the associated project accession numbers are given in table 2.

**Figure 1.**
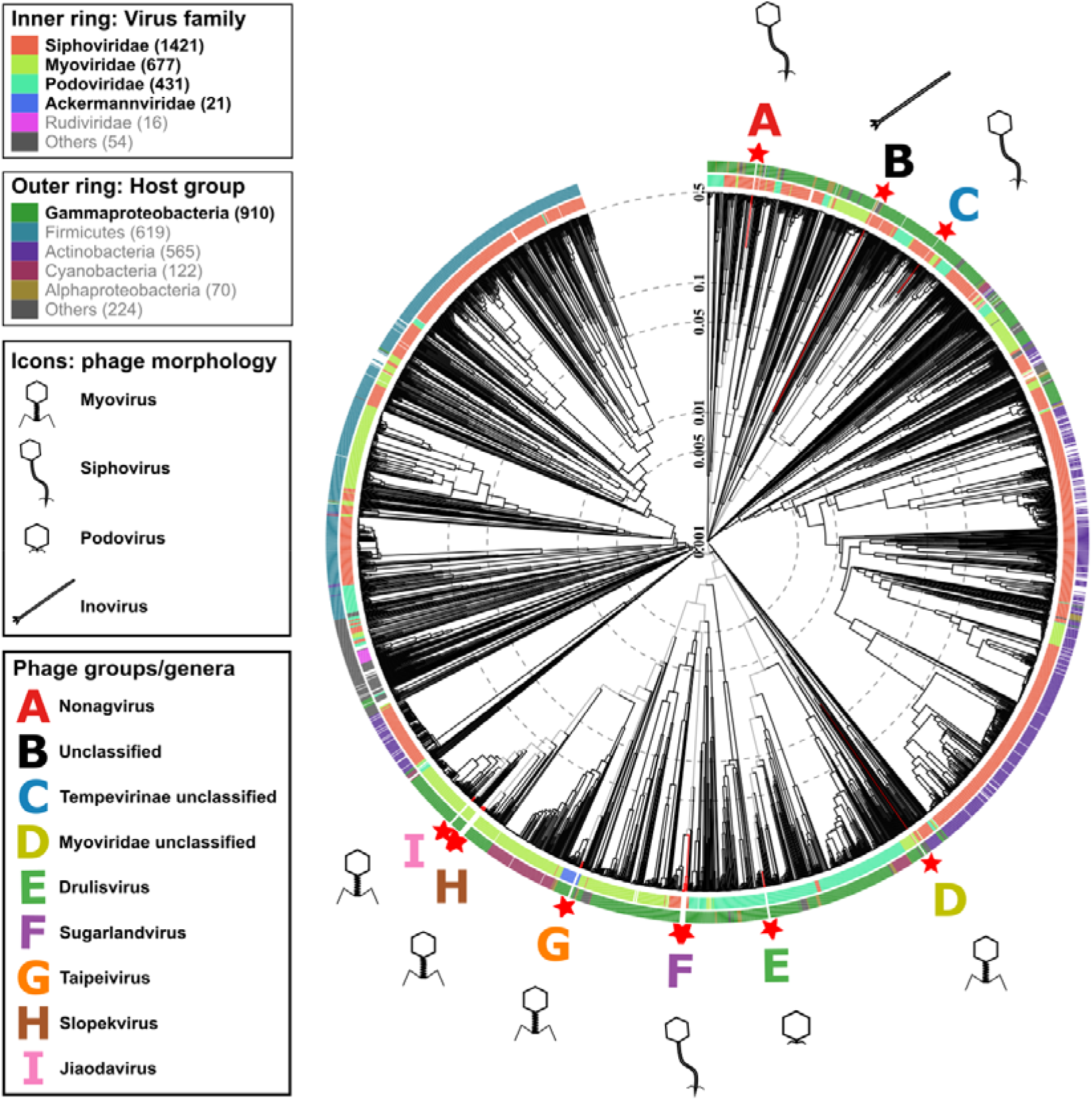
Protein level phylogenetic tree, generated by VIPtree. *Klebsiella* phage isolates (□). A-I denote phage groups with the following genera: A. *Nonagvirus;* B. unclassified family/genus; C. *Tempevirinae* unclassified; D. *Myoviridae* unclassified; E. *Drulisvirus;* F. *Sugarlandvirus;* G. *Taipeivirus;* H. *Slopekvirus* and I. *Jiaodavirus*. Icons indicate phage morphology.

**Figure 2.**
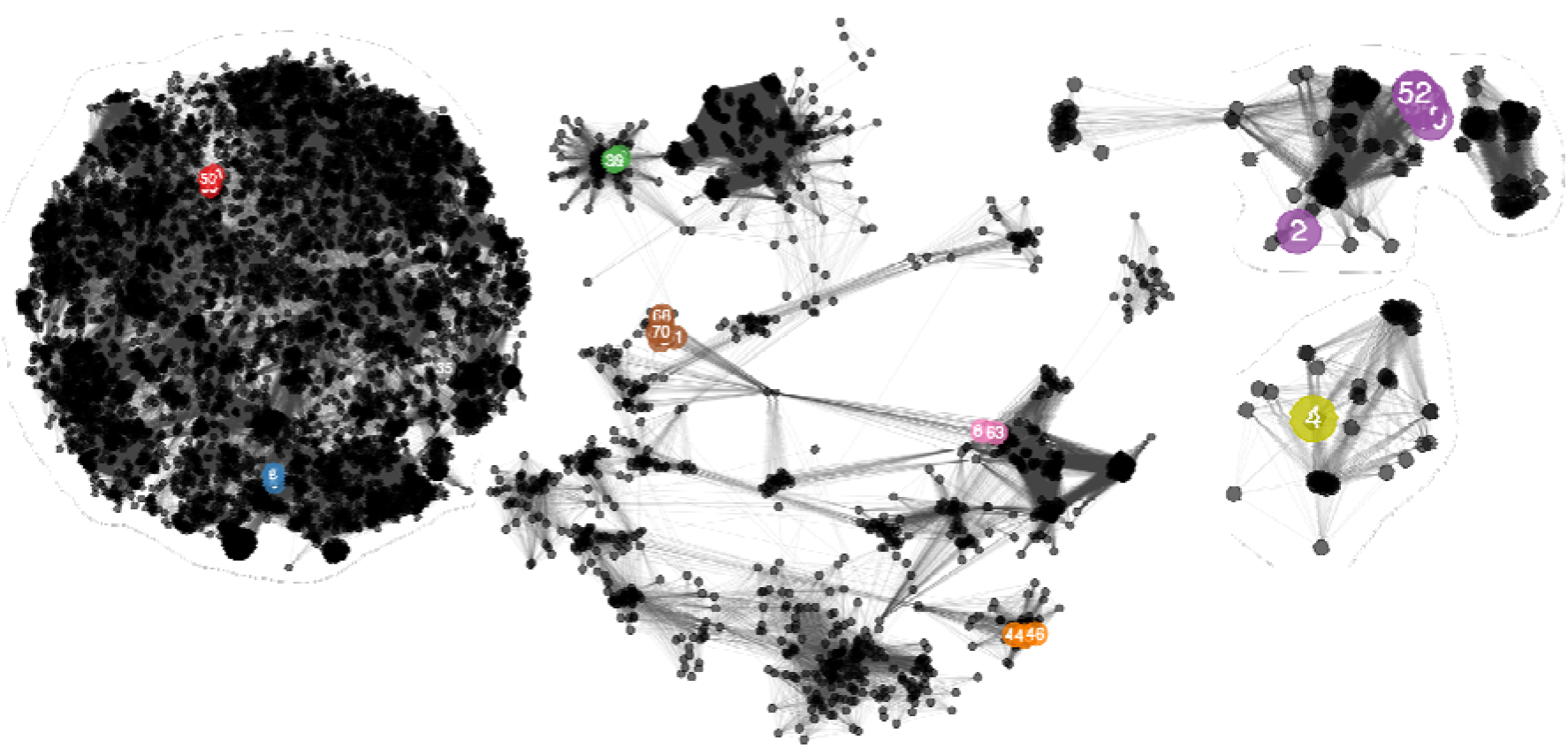
Network analysis of phage-encoded proteins calculated with vConTACT2. Coloured, numbered nodes represent our Klebsiella phage isolates, coloured according to the phage group and subsequent genera to which each phage belongs. Numbers within nodes indicate the lab identification numbers (see table 2). Smaller, black nodes represent previously sequenced phages as references. Edges between nodes represent shared proteins, such that many connecting edges implies greater pairwise shared protein content. Phage nodes are clustered based on shared proteins, with a spring-embedded (force directed) layout visualisation created in Cytoscape.

**Table 3.** Phage taxonomy and similarity to closest sequenced phage. Family, Subfamily and Genus are assigned based on the clustering patterns observed in vConTACT2 analysis (fig. 2), and conserved branching patterns observed in the marker gene phylogenetic trees (fig. S12-20). ANI refers to Average Nucleotide Identity calculated with orthoANI (https://www.ezbiocloud.net/tools/ani). Details for “comparison phage” relate to details of previously sequenced phages available in public databases used for ANI analysis.

| Phage name | Lab ID | Phage group | Taxonomy |  |  | Comparison phage |  |  |
| --- | --- | --- | --- | --- | --- | --- | --- | --- |
|  |  |  | Family | Subfamily | Genus | Phage name | Accession | ANI |
| vB_KppS-Raw | 33 | A | *Siphoviridae* |  | *Nonagvirus* | Enterobacteria phage JenP2 | KP719132 | 66.77 |
| vB_KppS-Eggy | 49 | A | *Siphoviridae* |  | *Nonagvirus* | Enterobacteria phage JenP2 | KP719132 | 65.86 |
| vB_KppS-Pokey | 50 | A | *Siphoviridae* |  | *Nonagvirus* | Enterobacteria phage JenP2 | KP719132 | 66.69 |
| vB_KppS-Ant | 35 | B | Unclassified |  |  | Caudovirales_phage_clone_3F_1 | MF417951 | 90.14 |
| vB_KaS-Gatomon | 6 | C | *Drexelviriidae* | *Tempevirinae* | unclassified | Escherichia phage Henu7 | MN019128 | 93.55 |
| vB_KaS-Ahsoka | 7 | C | *Drexelviriidae* | *Tempevirinae* | unclassified | Escherichia phage Henu7 | MN019128 | 92.09 |
| vB_KppS-Samwise | 8 | C | *Drexelviriidae* | *Tempevirinae* | unclassified | Escherichia phage Henu7 | MN019128 | 93.55 |
| vB_KvM-Eowyn | 4 | D | *Myoviridae* | unclassified |  | Serratia_phage_KpHz_2 | KF806589 | 95.02 |
| vB_KpP-Yoda | 43 | E | *Autographiviridae* | *Slopekvirinae* | *Drulivirus* | Klebsiella phage vB_KpnP_SU552A | KP708986 | 87.03 |
| vB_KqP-Goliath | 44 | E | *Autographiviridae* | *Slopekvirinae* | *Drulivirus* | Klebsiella phage vB_KpnP_SU552A | KP708986 | 85.82 |
| vB_KpP-Screen | 46 | E | *Autographiviridae* | *Slopekvirinae* | *Drulivirus* | Klebsiella phage vB_KpnP_SU552A | KP708986 | 86.58 |
| vB_KaS-Benoit | 1 | F | *Demereciviridae* |  | *Sugarlandvirus* | vB_Kpn_ME260 | NC_041899 | 94.29 |
| vB_KaS-Veronica | 2 | F | *Demereciviridae* |  | *Sugarlandvirus* | Klebsiella phage Sugarland | NC_042093 | 93.46 |
| vB_KppS-Totoro | 10 | F | *Demereciviridae* |  | *Sugarlandvirus* | vB_Kpn_ME260 | NC_041899 | 94.29 |
| vB_KppS-Ponyo | 19 | F | *Demereciviridae* |  | *Sugarlandvirus* | Klebsiella phage Sugarland | NC_042093 | 94.15 |
| vB_KppS-Jiji | 27 | F | *Demereciviridae* |  | *Sugarlandvirus* | Klebsiella phage Sugarland | NC_042093 | 93.26 |
| vB_KppS-Storm | 34 | F | *Demereciviridae* |  | *Sugarlandvirus* | Klebsiella phage Sugarland | NC_042093 | 95.02 |
| vB_KppS-Anoxic | 52 | F | *Demereciviridae* |  | *Sugarlandvirus* | vB_Kpn_ME260 | NC_041899 | 96.11 |
| vB_KqM-LilBean | 36 | G | *Ackermannviridae* |  | *Taipeivirus* | Klebsiella virus 0507KN21 | NC_022343 | 97.83 |
| vB_KqM-Bilbo | 38 | G | *Ackermannviridae* |  | *Taipeivirus* | Klebsiella virus 0507KN21 | NC_022343 | 97.22 |
| vB_KqM-Westerburg | 39 | G | *Ackermannviridae* |  | *Taipeivirus* | Klebsiella virus 0507KN21 | NC_022343 | 96.93 |
| vB_KoM-Pickle | 12 | H | *Myoviridae* | *Tevenvirinae* | *Slopekvirus* | Klebsiella phage KP15 | GU295964 | 97.93 |
| vB_KoM-Liquor | 61 | H | *Myoviridae* | *Tevenvirinae* | *Slopekvirus* | Klebsiella phage KP15 | GU295964 | 97.83 |
| vB_KpM-Milk | 62 | H | *Myoviridae* | *Tevenvirinae* | *Slopekvirus* | Klebsiella phage KP15 | GU295964 | 98.38 |
| vB_KpM-Mild | 65 | H | *Myoviridae* | *Tevenvirinae* | *Slopekvirus* | Klebsiella phage KP15 | GU295964 | 97.65 |
| vB_KpM-KalD | 67 | H | *Myoviridae* | *Tevenvirinae* | *Slopekvirus* | Klebsiella phage KP15 | GU295964 | 97.42 |
| vB_KoM-MeTiny | 68 | H | *Myoviridae* | *Tevenvirinae* | *Slopekvirus* | Klebsiella phage KP15 | GU295964 | 97.92 |
| vB_KpM-SoFaint | 70 | H | *Myoviridae* | *Tevenvirinae* | *Slopekvirus* | Klebsiella phage KP15 | GU295964 | 97.72 |
| vB_KoM-Flushed | 63 | I | *Myoviridae* | *Tevenvirinae* | *Jiaodavirus* | Klebsiella phage JD18 | KT239446 | 96.19 |
| vB_KpM-Wobble | 64 | I | *Myoviridae* | *Tevenvirinae* | *Jiaodavirus* | Klebsiella phage JD18 | KT239446 | 97.01 |

Alignments constructed in VIPtree^33^ showed varying levels of amino acid sequence identity but high gene synteny between our isolates and known phages (figures S3-10). The phage groups with the highest levels of amino acid identity and gene synteny to known phages were F, G, H and I corresponding to *Drulisvirus*, *Sugarlandvirus, Taipeivirus* and *Jiaodavirus* genera respectively (fig. S8-11). Phage isolates in group F, *Sugarlandvirus*, showed the greatest similarity to previously described phages (fig. S8). All *Sugarlandvirus* isolates grouped with previously described vB_Kpn_IME260 and *Klebsiella* phage Sugarland, except vB_KaS-Veronica which represents a new distinct species based on ANI. The *Sugarlandvirus* isolates did exhibit variation in their tail fibre genes (figure S9; at ~75 kb). In contrast, phage isolates in group A, corresponding to the known genus of *Nonagvirus*, were less similar to previously known phages (fig. S4).

There was low sequence identity, but high gene synteny between our isolates and previously sequenced phages for groups B, C and D (fig. S5-7), a result of the isolates in these groups have unresolved taxonomies. Interestingly, analysis of the phage with the smallest genome, vB_KppS-Ant of group B, revealed >99 % nucleotide identity to region of the *K. pneumoniae* 30104 genome (data not shown).

For further similarity analysis between our isolates and known phages, the protein sequences of three marker genes (DNA polymerase, major capsid protein and terminase large subunit) were used to draw phylogenetic trees (fig. S12-20). Conserved branching patterns, indicating close evolutionary history were observed between most of our phage isolates and known phages, confirming that our isolates from groups A, E, F, G, H and I belong to known genera (fig. S12,16-20), while groups B, C and D (fig. S13-15) do not. Given more distant relationships between the marker protein sequences, it is proposed that group C represent a novel genus of the subfamily *Tempevirinae* (fig. S14) and group D represent a novel genus of the *Myoviridae* family (fig. S15), while the marker genes of the phage isolate of group B had an even more distant relationship with phages of both *Siphoviridae* and *Myoviridae* families (fig. S13). Given the inovirus morphology observed for group B (fig. S2, black box), classification at the family level also remains unresolved.

### Host Range Testing

Most phages had a host range which extended past their original isolation host and was not explained by depolymerase activity. The number of strains infected by each phage is displayed in figure 3, further information on host range including strain and capsule type specificity is found in figure S1. Two of the isolated phage genera showed activity against only their isolation host, these are both comprised of putative temperate phages; group A (*Nonagvirus*; except KppS-Raw, which infected one additional strain) and group B (unclassified family/genera). The lytic phage groups D, E and G (*Myoviridae* unclassified, *Drulisvirus* and *Taipeivirus*, respectively) showed lytic activity against 3-7 *Klebsiella* strains and formed plaques in 1-7 of those strains. While group F (*Sugarlandvirus*) phages had activity against 16 *Klebsiella* strains and plaques on 10 strains. Group C (*Tempevirinae* unclassified) was highly variable, with KaS-Gatomon and KaS-Ahsoka forming plaques on only their isolation strain, while Kpp-Samwise formed plaques in seven strains and activity against a further five strains. The broadest range were lytic phages belonging to the subfamily *Tevenvirinae*; assigned to the groups H (*Slopekvirus*) and I (*Jiaodavirus*), demonstrated activity against 23 and 17 strains respectively and were able to produce plaques on 13 and 9 strains respectively. Within the genera host range varied between phages; from the group H (*Slopekvirus*), the phage KoM-Pickle was only able to form plaques on its isolation host, yet showed some level of activity against 22 out of 23 other *Klebsiella* spp. (figure 3; figure S1).

**Figure 3.**
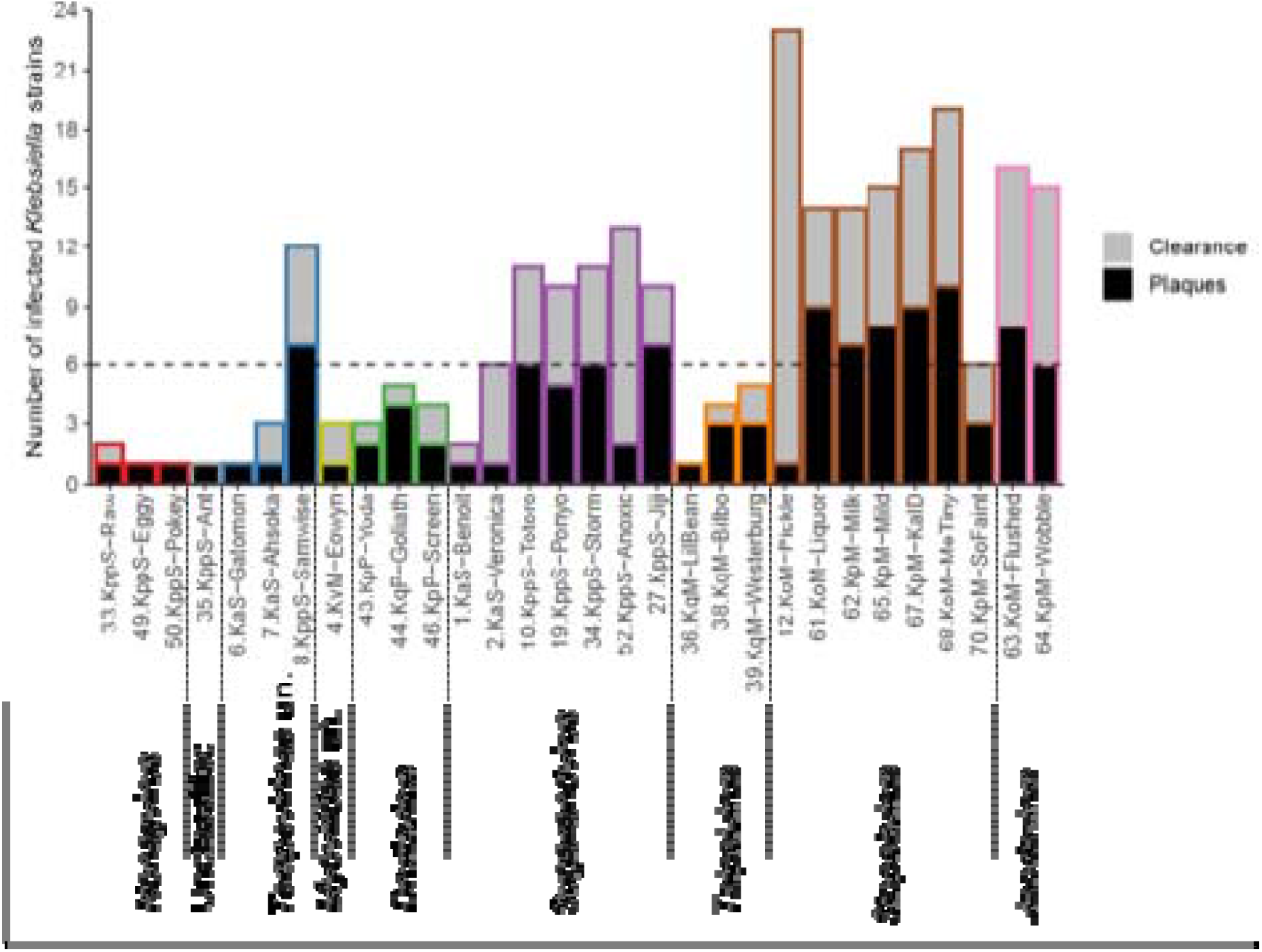
Number of *Klebsiella* strains infected by each bacteriophage out of a possible 24. Plaques, indicates the number of strains a phage replicated in and resulted in plaques on overlay agar; Clearance, indicates the additional number of strains the phage cleared or partially cleared the bacterial lawn during spot testing analysis (Figure S1). Totals include the isolation host. Bars marked with # denote an incomplete data set. Coloured outlines relate to the phage groups, as identified in figure 1, with genera annotated for each group.

### Phage Annotation

Genome annotation is notoriously difficult with phages, given the extremes of sequence variation evident in all phage proteins^50, 51^. As a first indication for comparisons, PROKKA was used and identified key phage genes e.g. portal proteins, capsid genes, tail proteins, components of the DNA replication and packaging machinery. Additionally, PhoH was a common feature in 18 of the 30 phages sequenced, including phages from groups D, F, G and H (*Myoviridae* unclassified, *Sugarlandvirus*, *Taipeivirus* and *Slopekvirus*, respectively). Holin and lysin pairs were identified in groups A, C and G (*Nonagvirus*, *Tempevirinae* unclassified and *Taipeivirus*, respectively), while endolysin and Rz1 spanin complex genes were identified in groups E and I (*Drulisvirus* and *Jiaodavirus*, respectively).

Most of our isolated phages encode at least one gene annotated as a putative “tail-fibre” or “tail-spike” protein (table S2). Intial structural predictions suggested that these proteins adopt beta-helical structures, a common protein architecture of both tail-spike proteins and the capsule depolymerase enzymes that are suggested to have evolved from these purely structural proteins^52–54^. These proteins also contained predicted enzymatic domains, such as the Pectate_lyase_3 domain or Peptidase_S74 domain, which have been identified in other phage encoded depolymerases^55–58^. Several of the analysed phage groups: B, C, F, H and I, did not have a predicted tail-fiber depolymerase protein. It is unclear whether these phages genuinely lack the activity, or that detection was hampered by extremes of sequence variation^50, 51^.

For those phages where a candidate depolymerase could be identifed, protein sequence relationships were mapped on a tree (figure 4). Phages in the group A (*Nonagvirus*), and group E (*Drulisvirus*) encode a similar type of putative predicted depolymerase protein. The putative depolymerases from *Drulisvirus* phages share high sequence conservation (~95 % identity, 100 % query) to the experimentally characterised depolymerase Kpv74_56 from the closely related *Drulisvirus*, *Klebsiella pneumoniae* phage KpV74^59^, and both sets of phage infect K2 capsule-producing strains of *Klebsiella* (fig. 4). Interestingly, the phages of two further genera; group D (*Myoviridae* unclassified; vB_KqM-Eowyn) and group G (*Taipeivirus*; vB_KqvM-LilBean, vB_KqvM-Bilbo and vB_KqvM-Westerburg) each contain multiple putative depolymerase-like proteins (7, 4, 4, 5 respectively; fig. 4). The putative depolymerase-like proteins of vB_KqM-Eowyn (D, *Myoviridae* unclassified) show low identity to previously characterised depolymerases (fig. 4). From our host range assays these phages showed activity towards strains that produced a small subset of K-antigens (vB_KqM-Eowyn – KL16, KL110; vB_KqvM-LilBean – KL35; vB_KqvM-Bilbo – KL35, K2 and vB_KqvM-Westerburg - KL35, KL2, KL3).

**Figure 4.**
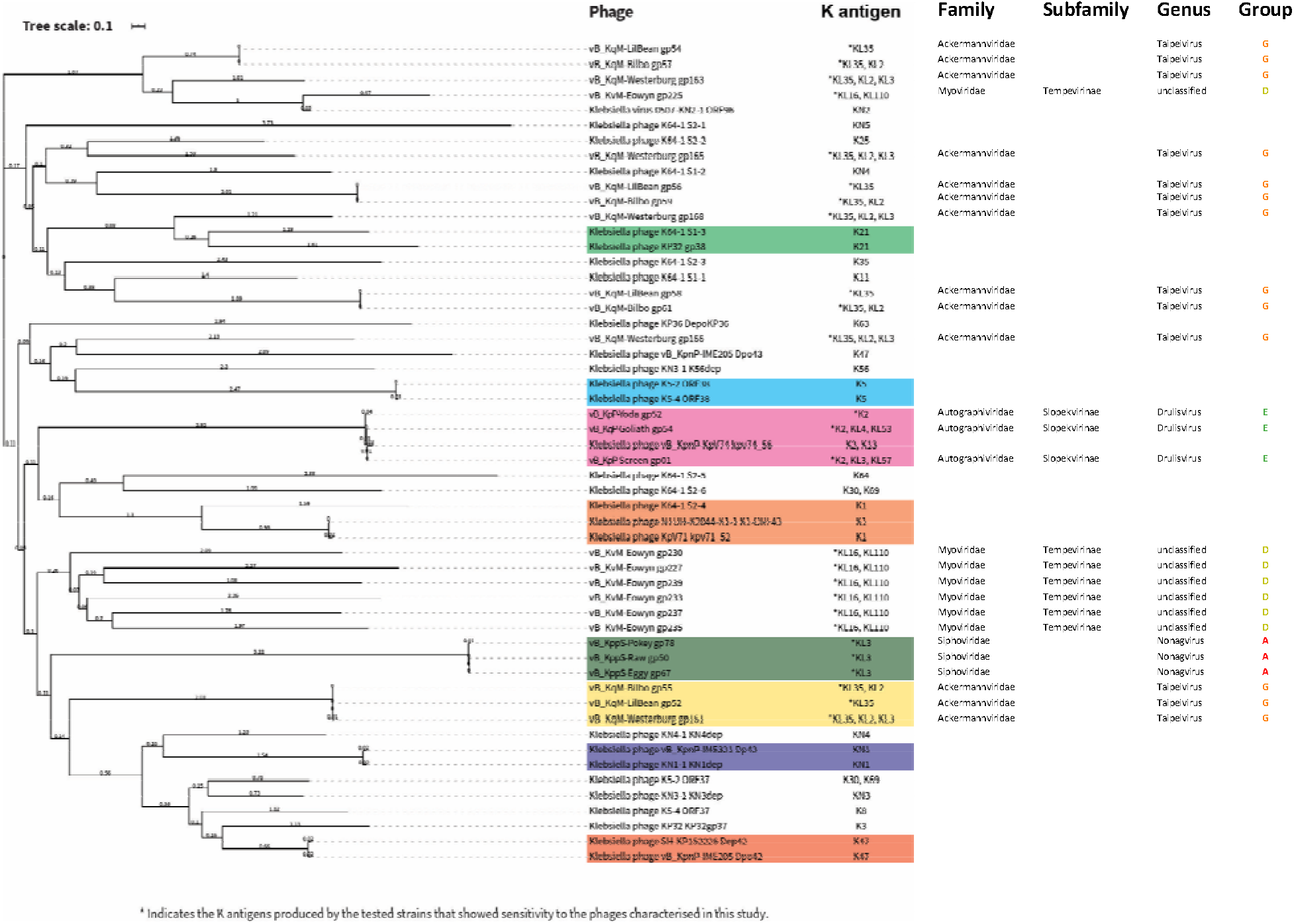
Phylogenetic tree of previously biochemically characterised depolymerase proteins that target *Klebsiella* spp. and putative depolymerase proteins in our phage isolates. Sequences were aligned with Muscle using SeaView. Phylogenetic tree construction was performed with MegaX with 500 bootstrap calculations using the LG model. Tree topology searches were performed using a combination of NNI and NJ/BioNJ. The tree was subsequently visualised and annotated using iTOL (v4). Depolymerases highlighted in colour blocks have overlapping potential target K antigens.

### Morphology of Phages

Phage-induced plaques in the lawns of host bacteria varied in size, and also varied in the presence or absence of halos surrounding the phage plaques. Representative images of phage plaques for each described genera are presented in figure 5 and images for all phages are included in supplementary figure S2. A diffuse halo around the phage plaques was observed in 26 of the 30 phage isolates (table S3 and fig. S2).

**Figure 5.**
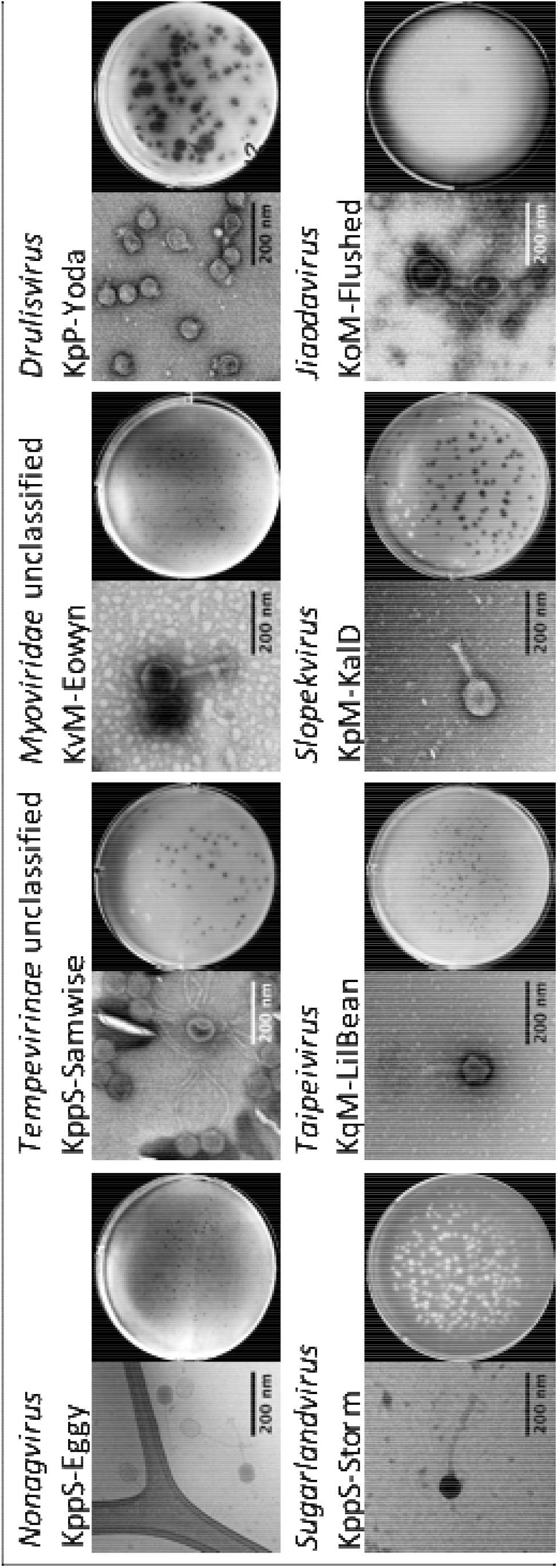
Phage morphology, showing representative TEM and plaque images for each phage group. TEM and plaque assays were performed as described in the methods. Scale bars on the TEM images represent 200 nm. Further images of all phages can be found in the supplementary materials, figure S2.

Representative TEM images of the described phage genera are provided in figure 5. Average tail length and capsid widths are given in supplementary table S3 based on 30 phage particles, a representative TEM image for each genera is displayed in figure S2. Of the phages imaged, 13 are myovirus, 13 are siphovirus, three are podovirus and one inovirus (fig. S2). The largest phage in this study was KvM-Eowyn of group D (*Myoviridae* unclassified), which had a capsid width of 140 nm and tail length of 140 nm, this corresponded to the largest genome at 269 Kbp. The smallest phage was KqP-Goliath, a podovirus of group E (*Drulisvirus*), with a capsid width of 41 nm and tail length of 10 nm, had the second smallest genome at 44 Kbp, after the filamentous prophage KppS-Ant, group B (unclassified family/genus). KppS-Raw, a group A (*Nonagvirus*) siphovirus, had a comparable capsid size of 46 nm, but a substantially longer tail (153 nm) and larger genome of 61 Kbp. The smallest icosahedral phage virions, group E (*Drulisvirus*), produced the largest plaques and halos, hence the name KqP-Goliath (table S1, figure S2).

### Phage Lysis period and Virulence in Host Strains

The lysis period and virulence indices (VP and MV50) of each phage, in their relevant isolation host strain in LB at 37 °C, are displayed in Figure 6. For six phages a lysis period was not achieved (KppS-Eggy, KppS-Pokey, KppS-Ant, KpM-Milk, KpM-KalD and KpM-SoFaint), the growth curve of the host bacteria was dampened (except KppS-Ant), but the culture density did not crash compared to the positive control - indicative of temperate phages. These phages also demonstrated a below average VP, close to 0 (with the exception of KpM-KalD), indicating little difference in the area under the curve between the control and phage infected cultures (fig. 6). Of the remaining phages that did exhibit a lysis period, the median time was ~70 minutes, ranging from 15 minutes to 210 minutes. There was no correlation between lysis period and the virulence measures (figure 6; statistical data not shown), implying that lysis period was not the driving factor. The average VP for the phages was 0.33 (range 0.06-0.64) and MV50 was achieved with a projected MOI of 4.17×10^17^ (range 3.5×10^-7^ - 1×10^19^), when the putative temperate phages from groups A and B were excluded. The putative temperate phages had lower virulence measures, reinforcing previous findings. For these, the average VP was 0.06 (range −0.07-0.36) and for the MV50 an MOI of 6.00×10^58^ (range 0.012-3×10^59^). Phages from groups C, E, F and I (*Tempevirinae* unclassified, *Drulisvirus*, *Sugarlandvirus*, *Jiaodavirus*, respectively) had the highest virulence (VP) (range 0.12-0.64, average, 0.42) and below average MV50 (range 3.5×10^-7^ - 1.30×10^5^, average 9.30×10^3^) indicated that they killed their hosts quickest, and needed a lower phage:host ratio to achieve this. It is however difficult to generalise for each genus because the two metrics varied between phages of the same genera. The virulence index results are specific to each phage and conditions assessed, and therefore are not directly comparable between phages grown on different hosts.

**Figure 6.**
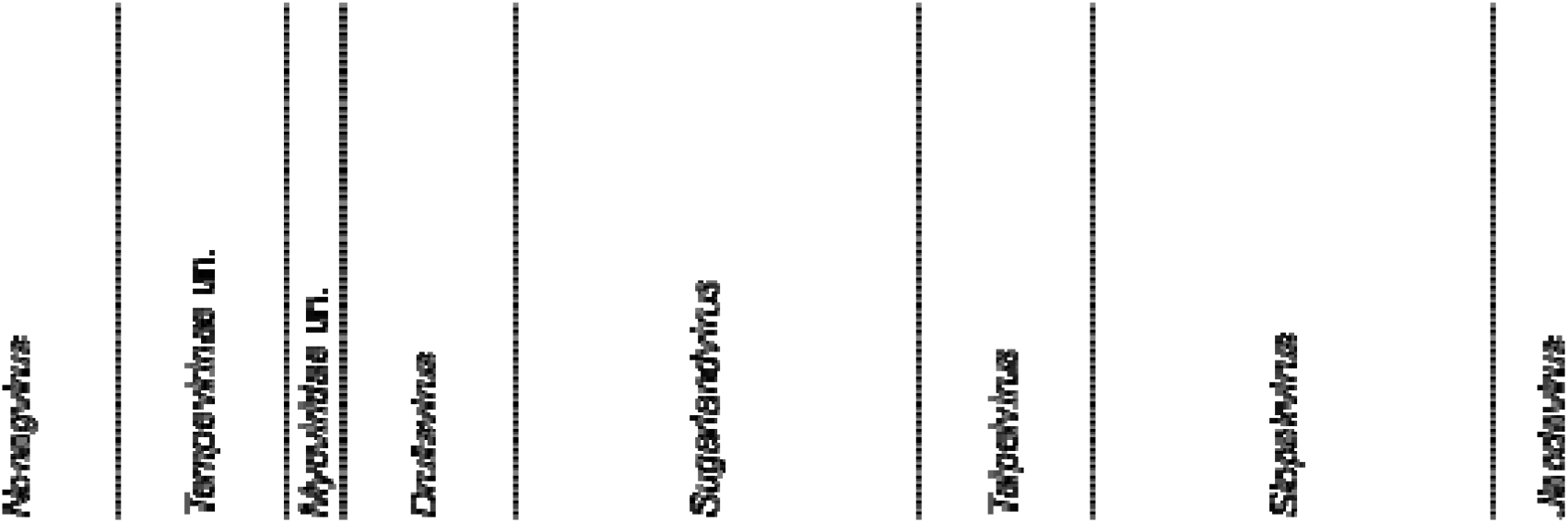
Lysis period and virulence indices are not correlated, but temperate phages are less virulent. Panel (A) displays the virulence index of the phages. Virulence index is a quantified measure of the phage, in their respective isolation host (table 2), in LB with 5 mM each of CaCl_2_ and MgCl_2_, at 37°C. Panel (B) displays the MV50, the MOI at which each phage achieves 50% of their maximal theoretical virulence. Both of these virulence measures are described in more detail by Storms^47^. Panel (C) displays the lysis period of the phage. Where this is left blank, a lysis period could not be established usually indicating temperate lifestyle. Coloured outlines relate to the phage groups, as identified in figure 1, with genera annotated for each group. Dashed lines indicate the median values for each metric.

## Discussion

*Klebsiella*-infecting phages, belonging to nine phylogenetically distinct lineages, were isolated from water samples sourced from different environments. The 30 independent phages were discovered using a panel of *Klebsiella* species - both clinical isolates and environmental strains - belonging to five bacterial species: *Klebsiella pnuemoniae, K. oxytoca, K. quasipneumoniae, K. aerogenes*, and *K. variicola*. The *Klebsiella* spp. used span 18 different capsule types.

This study discovered several phages and phage genera that have not previously been described. Phylogenetic analysis revealed that the filamentous phage vB_KppS-Ant of group B did not significantly cluster with any known phages at the shared protein-level, and therefore represents a novel genus. The genome reconstruction of this ssDNA phage is unexpected; our sequencing strategy was optimised for dsDNA, however previously studies have demonstrated Illumina sequencing to be inefficient yet successful at sequencing ssDNA phages^60^. The high sequence similarity of phage vB_KppS-Ant to a region of the *K. pneumoniae* 30104 genome (>99 %) indicates that it is an induced prophage, which represents a new genus. By combining shared protein network analysis and marker gene phylogenetic tree analysis, we identified two further novel phage genera: group C (*Tempevirinae* unclassified) and group D (*Myoviridae* unclassified).

### Factors determining host range

In general, comparative genomics revealed sequence conservation over a large portion of their genomes, with variability in only a few genes (fig. S4-11), with divergent characteristics in terms of host range and virulence. Despite several of these phage species having been identified in previous studies, there are examples where these showed differences in host-range. Thus, even within the small sample presented here, there is information to be gained about the factors determining host-range.

It has been suggested that selection pressure imposed from the use of a host is sufficient to amplify nongenetic variants of a phage that can cross host-range^47, 61^, and it also remains possible that uncharacterised genes, encoding proteins of unknown function, could adapt a given phage to a distinct host^62^. By way of example, the phages of group F (*Sugarlandvirus*) showed a high degree of similarity between genome sequences, with most variation concentrated in their tail fibre genes. Tail fibres mediate interaction with host cell receptors and are frequently rearranged in phages, allowing them to adhere to bacterial hosts^63^. These tail fibres can include domains with enzymatic function, enabling degradation of host-specific features such as polysaccharide capsules^64^. Three phages with >99% similarity in genome sequence were isolated on distinct hosts: *K. pneumoniae* 30104 (vB_KppS_Ponyo and vB_KppS_Totoro) or *K. aerogenes* 30053 (vB_KaS-Benoit). vB_KppS_Ponyo and vB_KaS-Benoit are 100 % identical at the nucleotide level, and vB_KppS_Totoro has a single SNP. Further host range analysis showed that the two phages propagated on *K. pneumoniae* 30104 had comparable host ranges, while the phage propagated on *K. aerogenes* 30053 had a different host range (fig. S1). This indicates that the propagation host, influenced host range, and that genome sequence cannot be used to infer host-range without careful consideration^65^. This is a vitally important point to consider when choosing phages for phage therapy, we cannot generalise based on genome similarity, when two phages of the same species and even those with 100% identity do not necessarily behave in the same way, particularly depending on prevailing conditions^66^.

While multiple features of strain-specific bacterial immunity can protect against phage replication in a given lineage, in *Klebsiella* the primary defense against both phages and antibiotics is a protective polysaccharide capsule^67, 68^. This capsule forms the outermost layer of the *Klebsiella* cell and acts as an important virulence factor^69^. There are currently at least 77 different serologically defined *Klebsiella* capsule types^70, 71^. Currently the use of whole genome methods for capsule typing is favoured^32^ and showed that our *Klebsiella* panel encompassed 18 capsule types, including three KL2 strains, an important capsule type in clinical infections and therefore of interest to develop effective therapies against^72^.

*Klebsiella* phages have repeatedly shown to be specific to host capsule types^73–75^, this is often linked to phage sugar-degrading enzymes called depolymerases that target specific capsule types^56, 59, 76^. Given the broad host ranges observed in our collection of phages (fig. 3) and depolymerase indicative halos^77^ in 83% of our phage isolates categorised as groups C-I (table S3), we sought to identify depolymerases genes. A surprisingly high number of putative depolymerase genes (7) were identified in the group D (*Myoviridae* unclassified) phage KvM-Eowyn, which only produced plaques in its KL16 host strain, but showed potential depolymerase activity against a KL110 producing strain. *Klebsiella* phages encoding up to 11 different polysaccharide depolymerase genes have previously been characterised, but have also been shown to infect a correspondingly wide range of *Klebsiella* capsule types^78^, indicating the need to expand our *Klebsiella* panel. The phages in group G (*Taipeivirus*) encoded 4-6 each depolymerases, these phages produced plaques against the clinically relevant KL2 capsule type. The phages of group A (*Nonagvirus*) and group E (*Drulisvirus*) had only one depolymerase gene each, in line the with the average number of expected depolymerases^53^. Characterisation of these putative depolymerase genes is important to further investigate the potential extended host range of these phages, beyond the *Klebsiella* strains included in this analysis. Conversely, genes encoding depolymerases were not identifiable in all of our halo producing phages, including in some of the broadest ranging phages of groups F, H and I (*Sugarlandvirus, Slopekvirus* and *Jiaodavirus*). The phage with the broadest host-range, KoM-MeTiny of group H (*Slopekvirus*), produced halos, showed depolymerase/lysis activity against 79% of the *Klebsiella* strains, and produced plaques on 42% of the *Klebsiella* tested, which included nine different capsule types, but had no identifiable depolymerase genes. We suggest that this is either because conventional depolymerases are not essential for the phages to permeate the capsule layer, or because of shortcomings in the sequence-based annotation of phage genomes^50, 51^. There is limited sequence conservation between many of the putative tail-fiber/tail-spike depolymerase proteins from our collection of phages to those that have been biochemically validated, therefore further characterisation of these proteins will be critical for future optimisation of phage cocktails for therapeutic uses.

### Application to future phage-based therapy

As expected with lytic phages, the phages isolated caused crashes in the *Klebsiella* cultures, but no phage was able to suppress *Klebsiella* growth for more than 12 hours (Figure S3). Phage cocktails are frequently used to improve the impact of phages on a *Klebsiella* population and are currently considered crucial for the efficacy of phage therapy^79–81^. Phage cocktails benefit from complementarity and redundancy between the combined phages to overcome host-evolved phage-resistance^82^, which may account for the resurgences seen in our *Klebsiella* cultures. After phage selection, the testing of phage combinations is imperative to ensure no adverse effects occur, such antagonistic phage interactions, which could result in bacterial stress responses or biofilm formation, as seen with sub-lethal antibiotic use^83, 84^. The phages described in this study have been supplied for use in compassionate phage therapy, requiring rigorous and lengthy testing to identify which phages were active against the clinically infective *Klebsiella* and to ensure safety.

For use in phage therapy, phages must not encode toxins or AMR genes^79, 81, 85^. None of our *Klebsiella* phages contained identifiable toxins or AMR genes. Other factors may be considerations for taking phages such as these into preclinical trials. For example, the host infection dynamics and low virulence of the phages of group A (*Nonagvirus*) indicate that these isolates may also be temperate and/or may not be curative on infections. The temperate phage KppS-Ant (group B), as the only phage with an identifiable integrase for lysogeny would be excluded for phage therapy purposes^86^. Additionally, the host infection dynamics and low virulence of the phages of group A (*Nonagvirus*) indicates a temperate lifestyle, paired with the knowledge that the closest related known phages are temperate, thus excluding these isolates for phage therapy. Group I (*Jiaodavirus*) phages encode a Hoc-like protein, which in phage T4 has been demonstrated to be highly immunogenic^87^. It should be established if these phages cause an immune response before using them for phage therapy.

The genomic information and experimental data presented here for phage groups C, F, G and H (*Tempevirinae* unclassified, *Sugarlandvirus, Taipeivirus* and *Slopekvirus*, respectively) indicates that they are lytic, not lysogenic, and thus suitable for preclinical evaluation. We suggest that the isolates of phage group F (*Sugarlandvirus*) and group H (*Slopekvirus*) are the best candidates for future development in phage therapy, given their broad host range, high virulence, short latency period and lack of potentially harmful genes. Taken together our data suggests that in order to provide universal, effective phage therapy against *Klebsiella* infections, a phage cocktail comprised of multiple diverse phages should be developed. This cocktail would prevent growth of *Klebsiella* cells resistant to a single phage or phage groups we isolated.

## Conclusions

A diverse range of *Klebsiella* phages were isolated from various environmental samples. The temperate phages discovered in this work may have interesting future applications, but are unsuitable for clinical applications. Phage isolates belonging to the *Sugarlandvirus* and *Slopekvirus* genera were deemed most suitable for phage therapy, due to their broad host range, high virulence, short lysis period and lack of potentially harmful genes. Despite some of our phage isolates grouping into a single previously described phage species, within species variation in both host range and virulence were observed. This demonstrates the necessity to microbiologically characterise phages before selecting candidates for therapeutic use.

## Acknowledgments

We acknowledge the help of Severn Trent for enabling us to collect sewage samples from Spernal sewage treatment works. We acknowledge the Midlands Regional Cryo-EM Facility, hosted at the Warwick Advanced Bioimaging Research Technology Platform, for use of the JEOL 2100Plus, supported by MRC award reference MC_PC_17136. Genome sequencing was provided by MicrobesNG (http://www.microbesng.uk). This work was supported by a Warwick Integrative Synthetic Biology (WISB) early career fellowship, funded jointly by BBSRC and EPSRC to EJ and the Monash Warwick Alliance Accelerator Fund October 2019 to EJ and TL. Bioinformatics analysis was carried out by infrastructure provided by MRC CLIMB (MR/T030062/1). The work has also been supported by PhD fellowships awarded to LK, LG, GM and HS funded by BBSRC/ESPRC DTPs.

---

Figure S1. Phage host range matrix. Host range was determined by spot testing on LB agar overlay plates against the *Klebsiella* spp. in our panel. Dark blue indicates host of isolation, in which plaques were produced, Light blue none-host strains where plaques were produced, Red indicates that spot testing caused the lawn to clear, but no plaques were visible, Dark red indicates some incomplete reduction was observed in the bacterial lawn, but no plaques and Grey indicates no observed effect.

Figure S2. Phage morphology; TEM and plaque morphology. The clearest TEM images were selected for each phage, scales vary for each image and nm scale bars are included for each image. An image of the plaques produced by each phage is included following overnight incubation at 37 °C on agar overlay plates.

Figure S3. Impact of each phage isolate on the growth curves of its reciprocal isolation host, grown in LB at 37 °C.

Figure S4. Group A (*Nonagvirus*) amino acid alignment of our phage isolates and reference genomes identified in vConTACT2 analysis, drawn in VIPtree.

Figure S5. Group B (unclassified family/genus) amino acid alignment of our phage isolates and reference genomes identified in vConTACT2 analysis, drawn in VIPtree.

Figure S6. Group C (*Tempevirinae* unclassified) amino acid alignment of our phage isolates and reference genomes identified in vConTACT2 analysis, drawn in VIPtree.

Figure S7. Group D (*Myoviridae* unclassified) amino acid alignment of our phage isolates and reference genomes identified in vConTACT2 analysis, drawn in VIPtree.

Figure S8. Group E (*Drulisvirus*) amino acid alignment of our phage isolates and reference genomes identified in vConTACT2 analysis, drawn in VIPtree.

Figure S9. Group F (*Sugarlandvirus*) amino acid alignment of our phage isolates and reference genomes identified in vConTACT2 analysis, drawn in VIPtree.

Figure S10. Group G (*Taipeivirus*) amino acid alignment of our phage isolates and reference genomes identified in vConTACT2 analysis, drawn in VIPtree.

Figure S11. Combined group H (*Slopekvirus*) and group I (*Jiaodavirus*) amino acid alignment of our phage isolates and reference genomes identified in vConTACT2 analysis, drawn in VIPtree.

Figure S12. Phylogenetic trees of DNA polymerase and terminase large subunit of phage Group A (*Nonagvirus*). Drawn with RaxML using the GAMMA model of heterogeneity and the maximumlikelihood method based on the JTT substitution matrix. Trees contain phage isolates from group A and reference phages identified by vConTACT2 and closest BLASTP hits. Coloured bar indicates known genera of reference phages.

Figure S13. Phylogenetic trees of terminase large subunit and major capsid protein of phage Group B (unclassified family/genus). Drawn with RaxML using the GAMMA model of heterogeneity and the maximum-likelihood method based on the JTT substitution matrix. Trees contain phage isolates from group B and reference phages identified by vConTACT2 and closest BLASTP hits. Coloured bar indicates known genera of reference phages.

Figure S14. Phylogenetic tree of terminase large subunit of phage Group C (*Tempevirinae* unclassified). Drawn with RaxML using the GAMMA model of heterogeneity and the maximumlikelihood method based on the JTT substitution matrix. Tree contains phage isolates from group C and reference phages identified by vConTACT2 and closest BLASTP hits. Coloured bar indicates known genera of reference phages.

Figure S15. Phylogenetic trees of DNA polymerase, major capsid protein and terminase large subunit of phage Group D (*Myoviridae* unclassified). Drawn with RaxML using the GAMMA model of heterogeneity and the maximum-likelihood method based on the JTT substitution matrix. Trees contain phage isolates from group D and reference phages identified by vConTACT2 and closest BLASTP hits. Coloured bar indicates known genera of reference phages.

Figure S16. Phylogenetic tree of DNA polymerase of phage Group E (*Drulisvirus*). Drawn with RaxML using the GAMMA model of heterogeneity and the maximum-likelihood method based on the JTT substitution matrix. Tree contains phage isolates from group E and reference phages identified by vConTACT2 and closest BLASTP hits. Coloured bar indicates known genera of reference phages.

Figure S17. Phylogenetic trees of DNA polymerase, major capsid protein and terminase large subunit of phage Group F (*Sugarlandvirus*). Drawn with RaxML using the GAMMA model of heterogeneity and the maximum-likelihood method based on the JTT substitution matrix. Trees contain phage isolates from group F and reference phages identified by vConTACT2 and closest BLASTP hits. Coloured bar indicates known genera of reference phages.

Figure S18. Phylogenetic trees of DNA polymerase, major capsid protein and terminase large subunit of phage Group G (*Taipeivirus*). Drawn with RaxML using the GAMMA model of heterogeneity and the maximum-likelihood method based on the JTT substitution matrix. Trees contain phage isolates from group G and reference phages identified by vConTACT2 and closest BLASTP hits. Coloured bar indicates known genera of reference phages.

Figure S19. Phylogenetic trees of DNA polymerase, major capsid protein and terminase large subunit of phage Group H (*Slopekvirus*). Drawn with RaxML using the GAMMA model of heterogeneity and the maximum-likelihood method based on the JTT substitution matrix. Trees contain phage isolates from group H and reference phages identified by vConTACT2 and closest BLASTP hits. Coloured bar indicates known genera of reference phages.

Figure S20. Phylogenetic trees of DNA polymerase, major capsid protein and terminase large subunit of phage Group I (*Jiaodavirus*). Drawn with RaxML using the GAMMA model of heterogeneity and the maximum-likelihood method based on the JTT substitution matrix. Trees contain phage isolates from group I and reference phages identified by vConTACT2 and closest BLASTP hits. Coloured bar indicates known genera of reference phages.

Table S1. List of characterised *Klebsiella* targeting depolymerase tail-fibre proteins used to construct the phylogenetic tree (Figure 4).

Table S2. List of putative depolymerase tail-fibre proteins from novel phages characterised in this study. Each sequence was analysed by BLASTP, HMMer and HHpred using the default parameters. The top HHpred hits are described for each protein. These sequences in combination with the sequences of characterised depolymerases from phages described in Table S1 were used to construct the phylogenetic tree (Figure 4).

Table S3. Phage measurements; phage particle measurement calculated from TEM imaging and measured with imageJ, and genome size in nucleotide base pairs.

