## Supplementary figures and images for "Isolation and characterisation of *Klebsiella* phages for phage therapy"

### Supplementarty fig S1

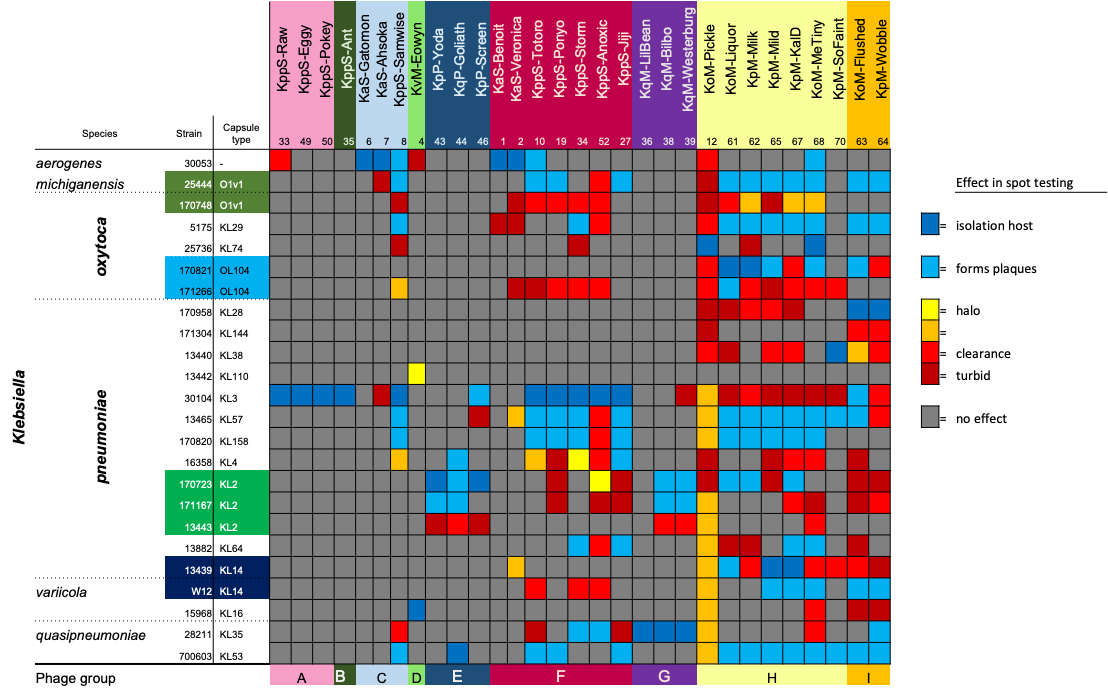

### Supplementarty fig S2

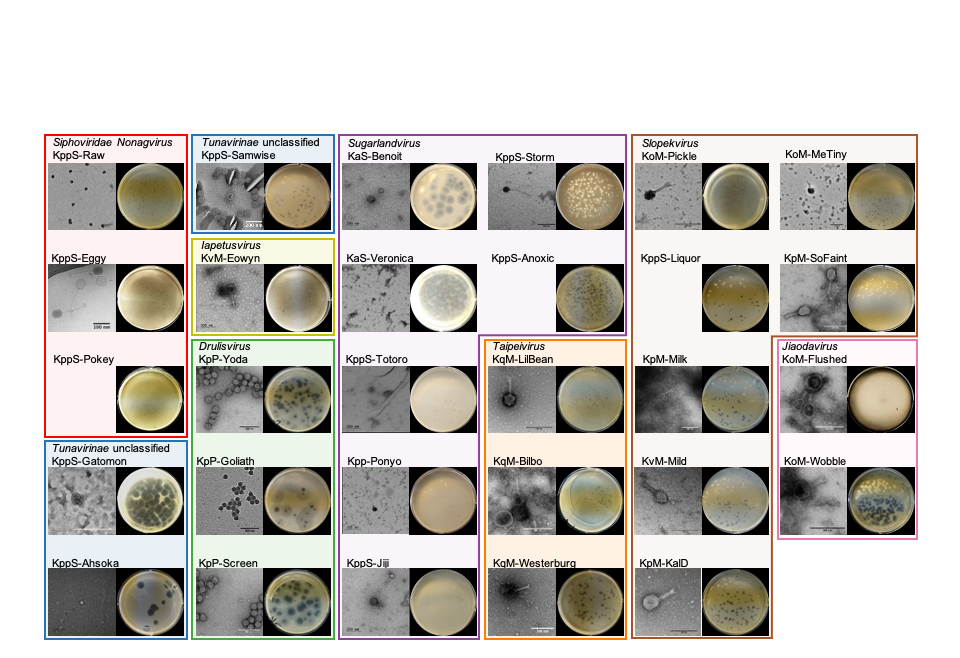

### Supplementarty fig S3

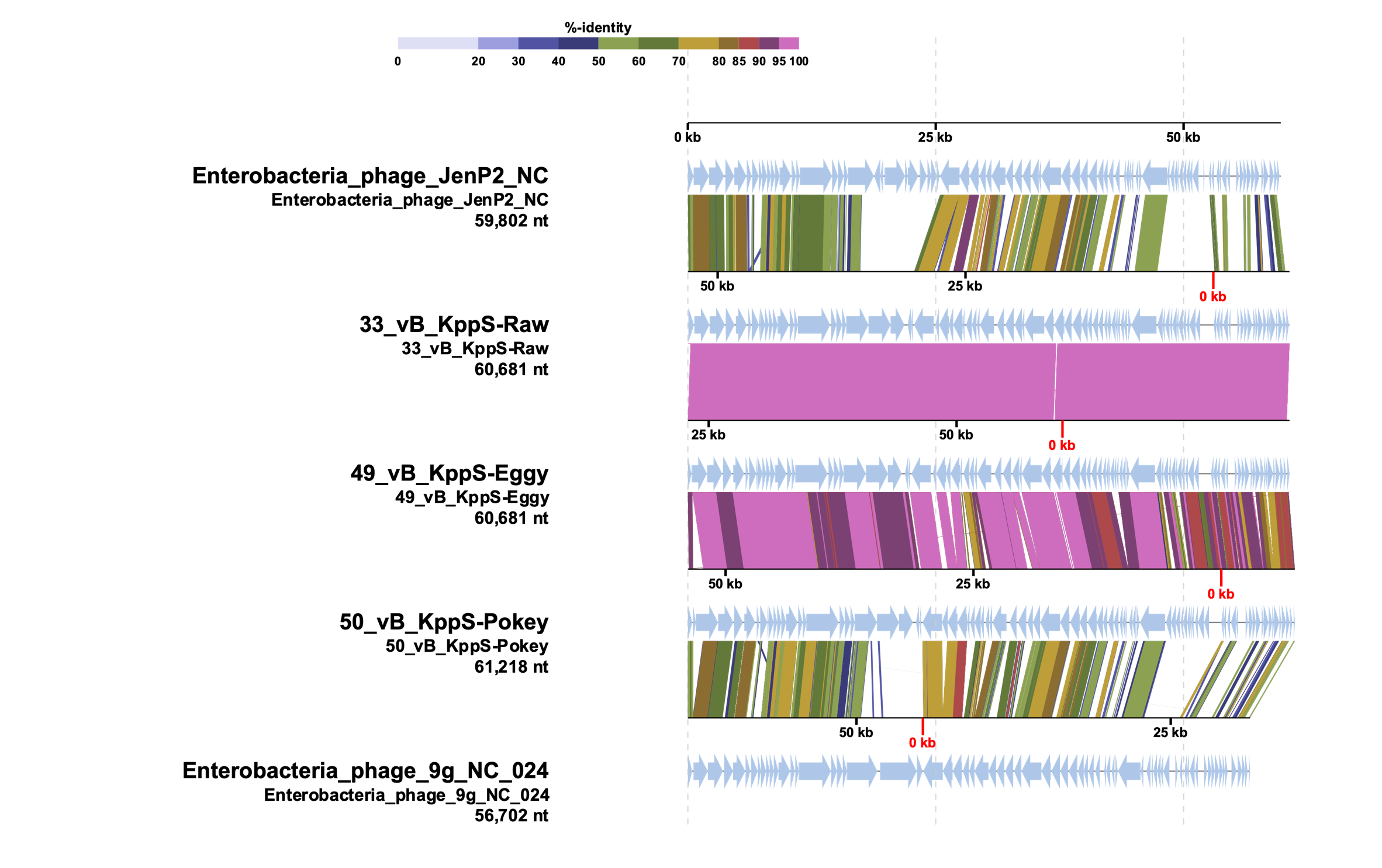

### Supplementarty fig S4

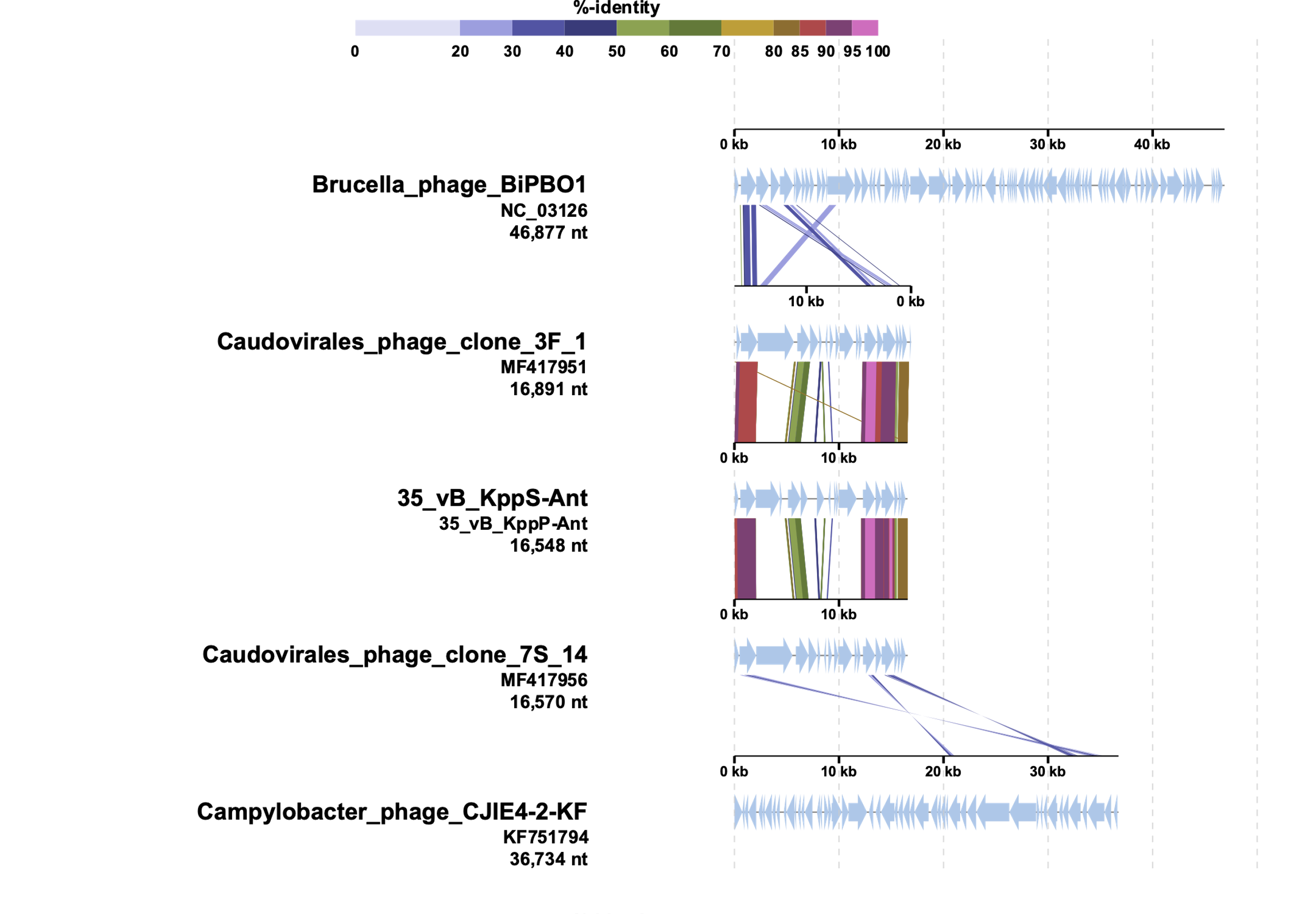

### Supplementarty fig S5

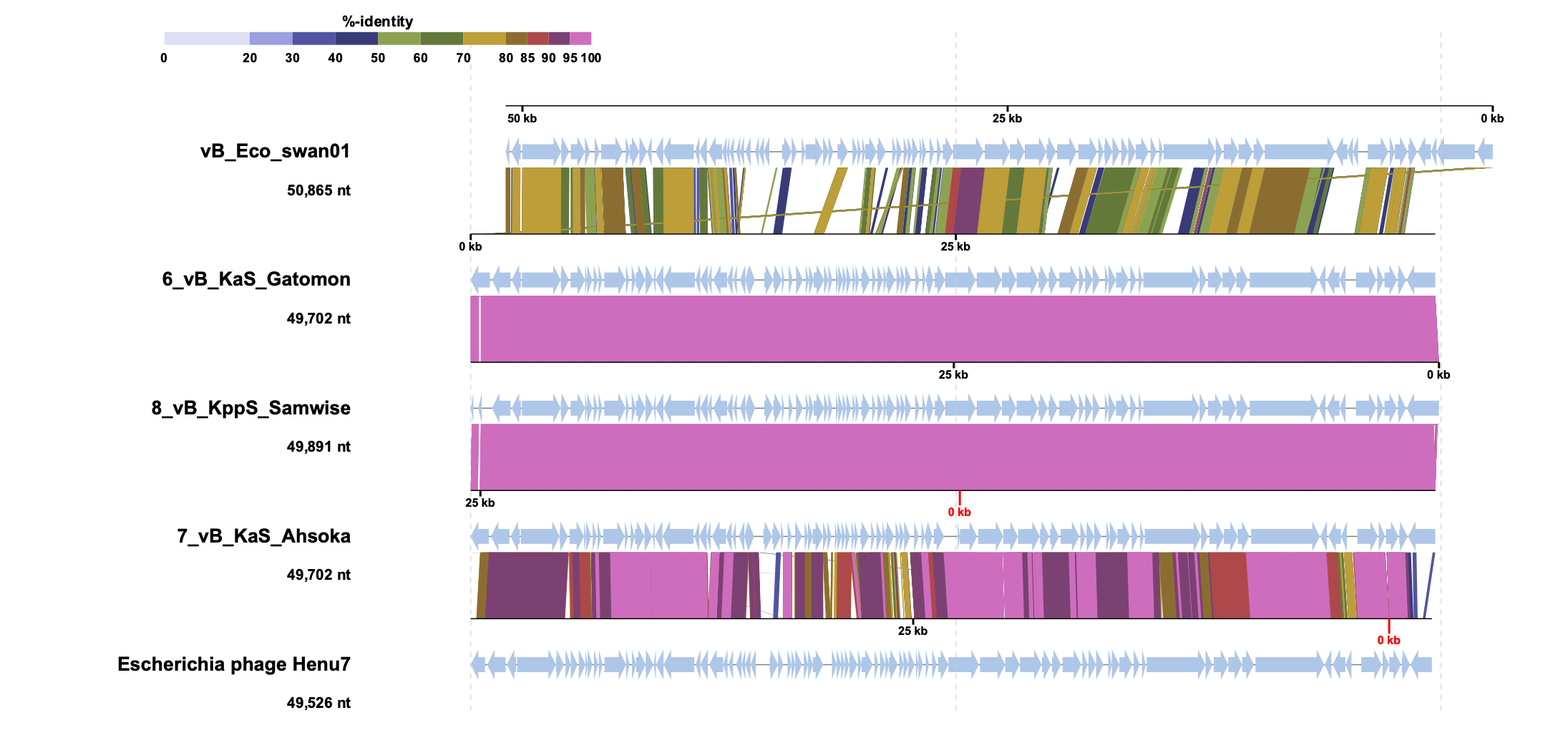

### Supplementarty fig S6

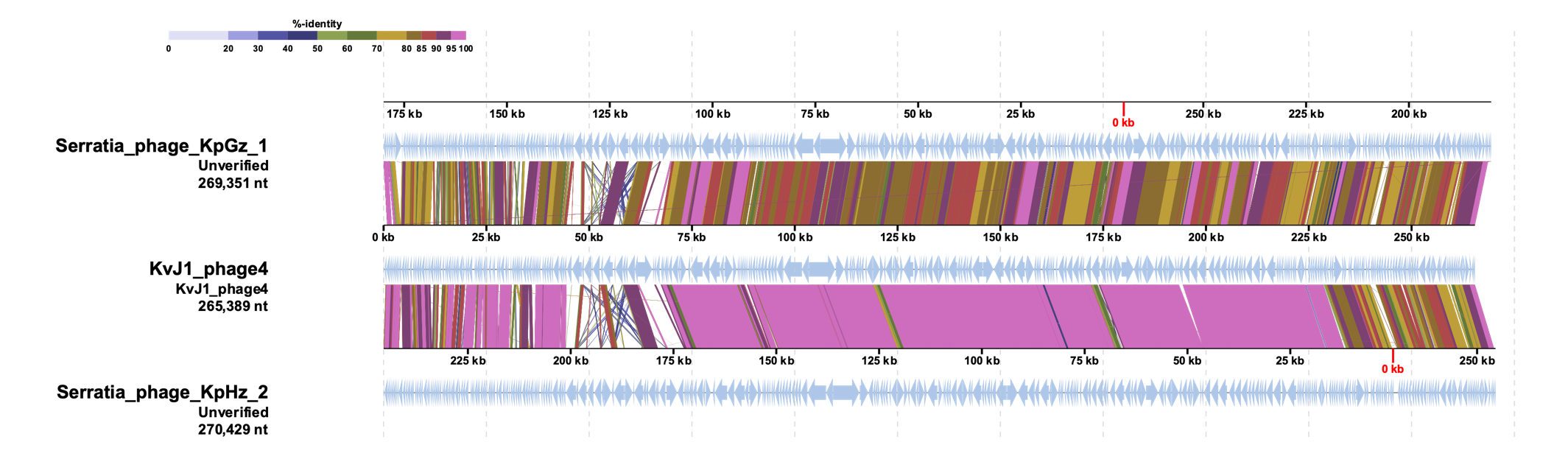

### Supplementarty fig S7

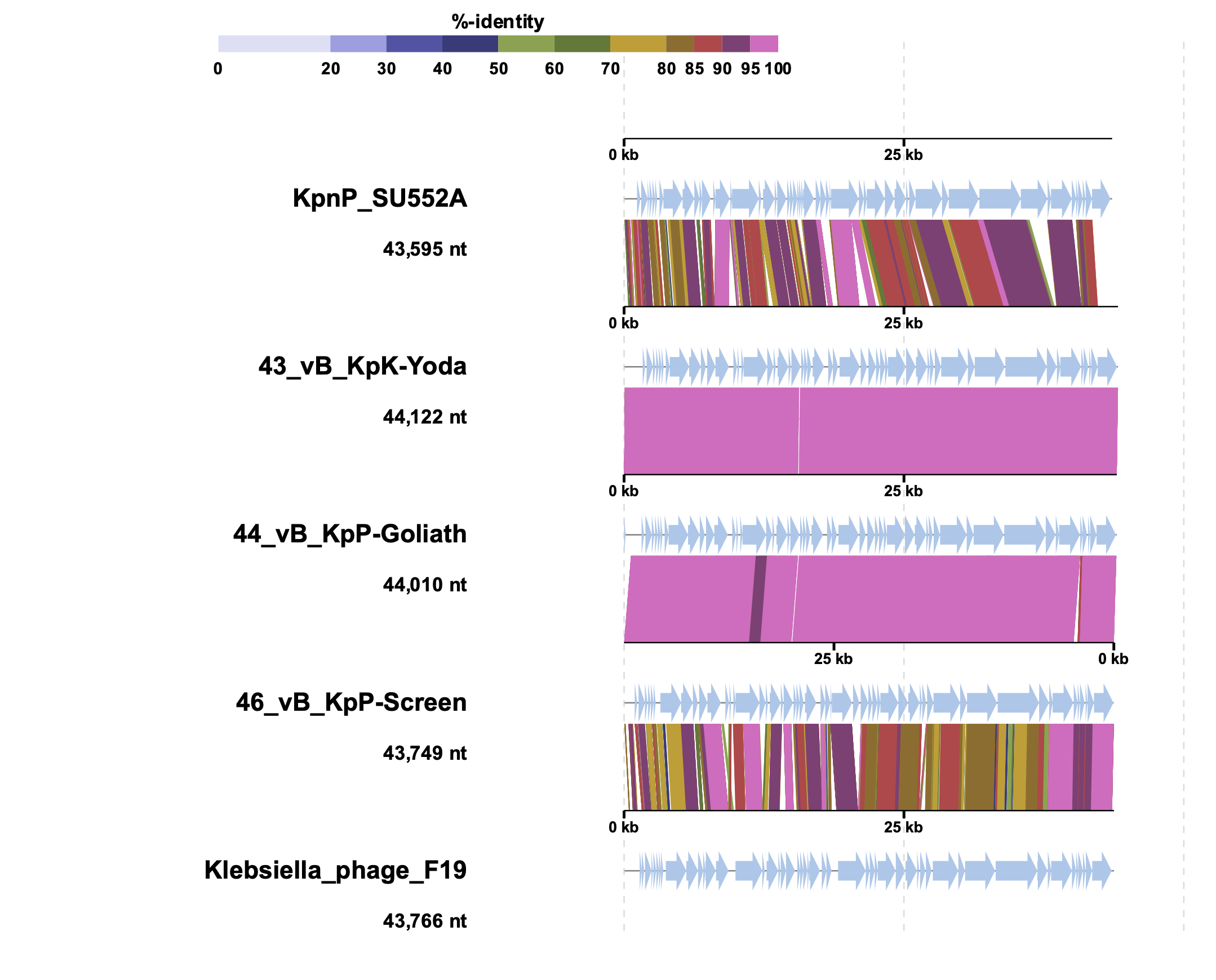

### Supplementarty fig S8

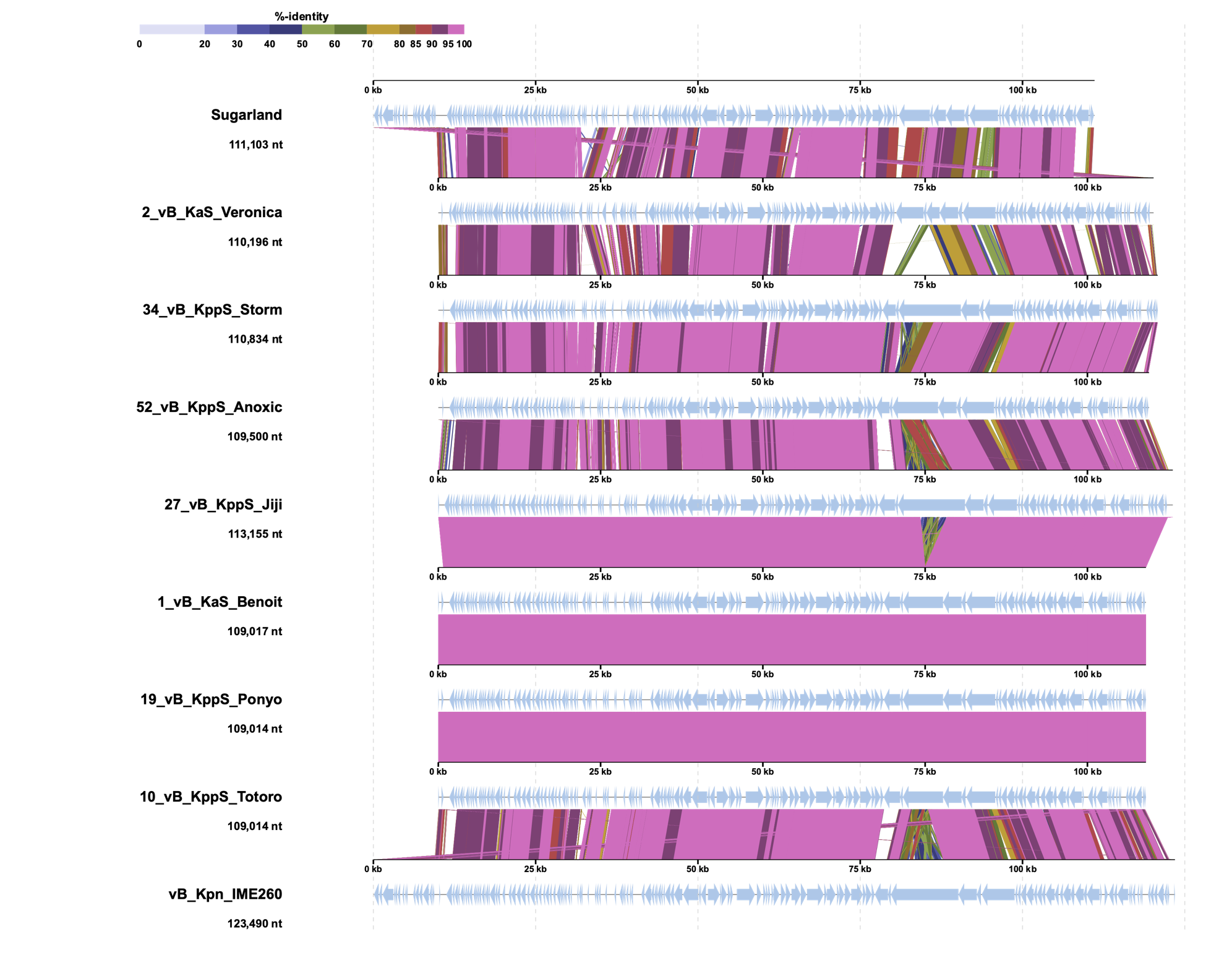

### Supplementarty fig S9

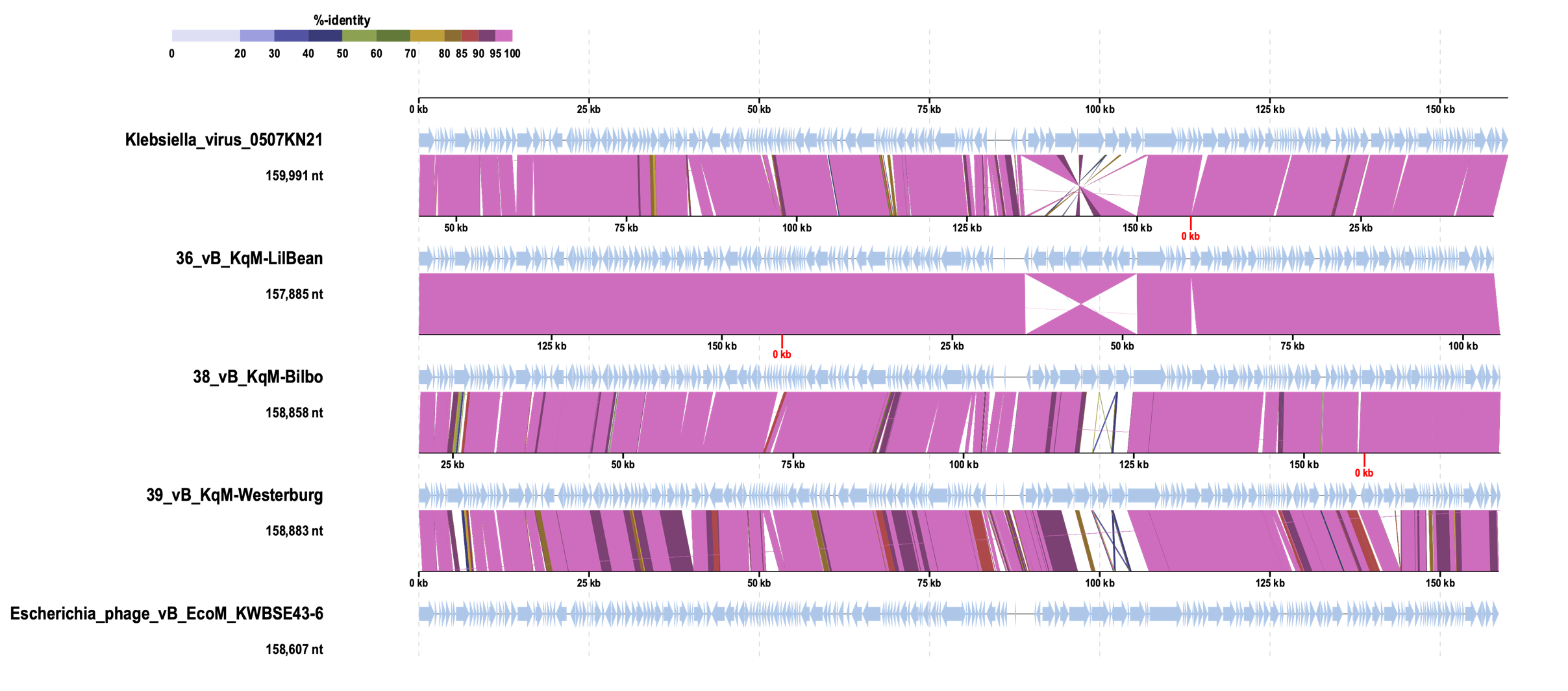

### Supplementarty fig S10

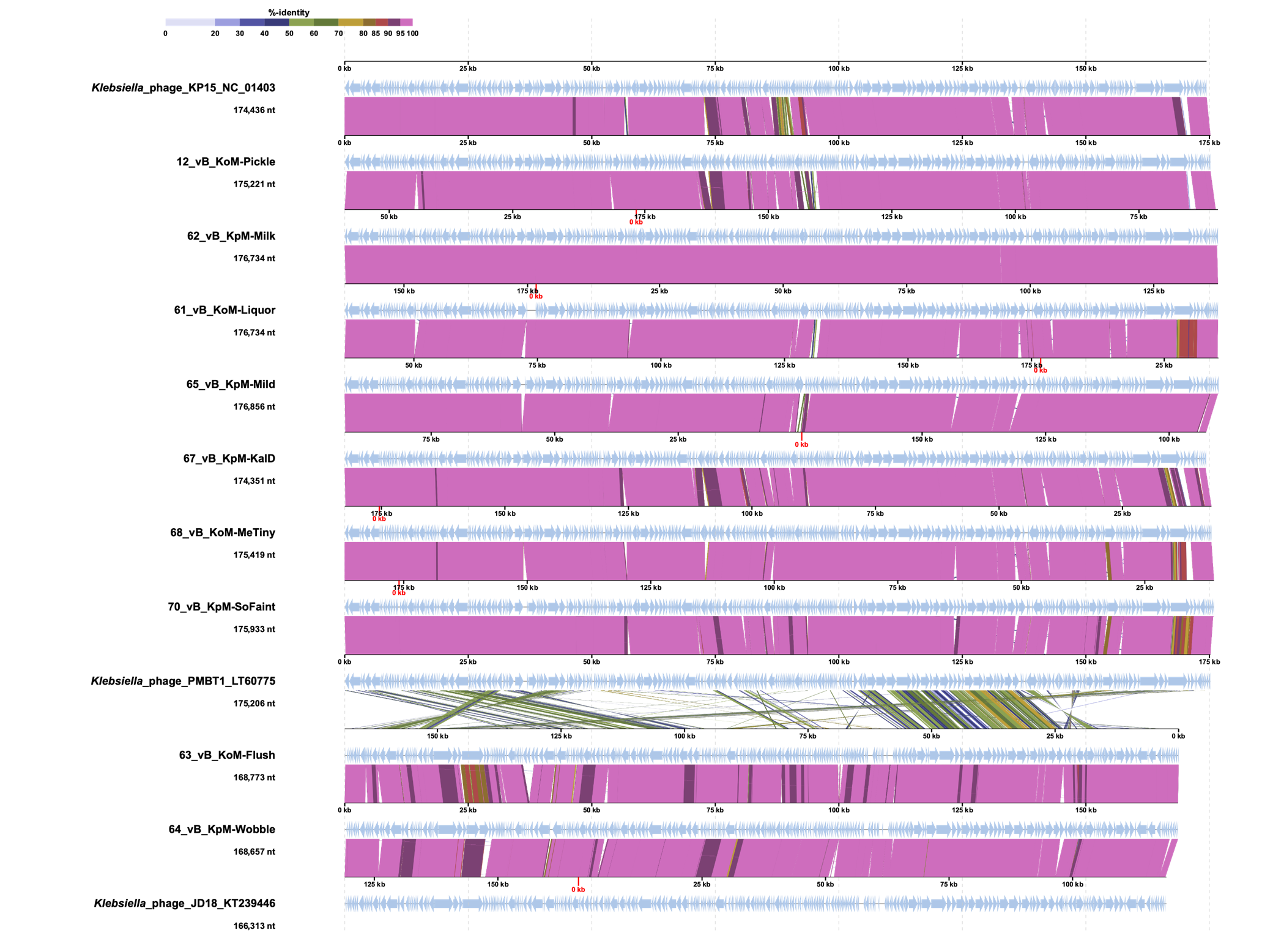

### Supplementarty fig S11

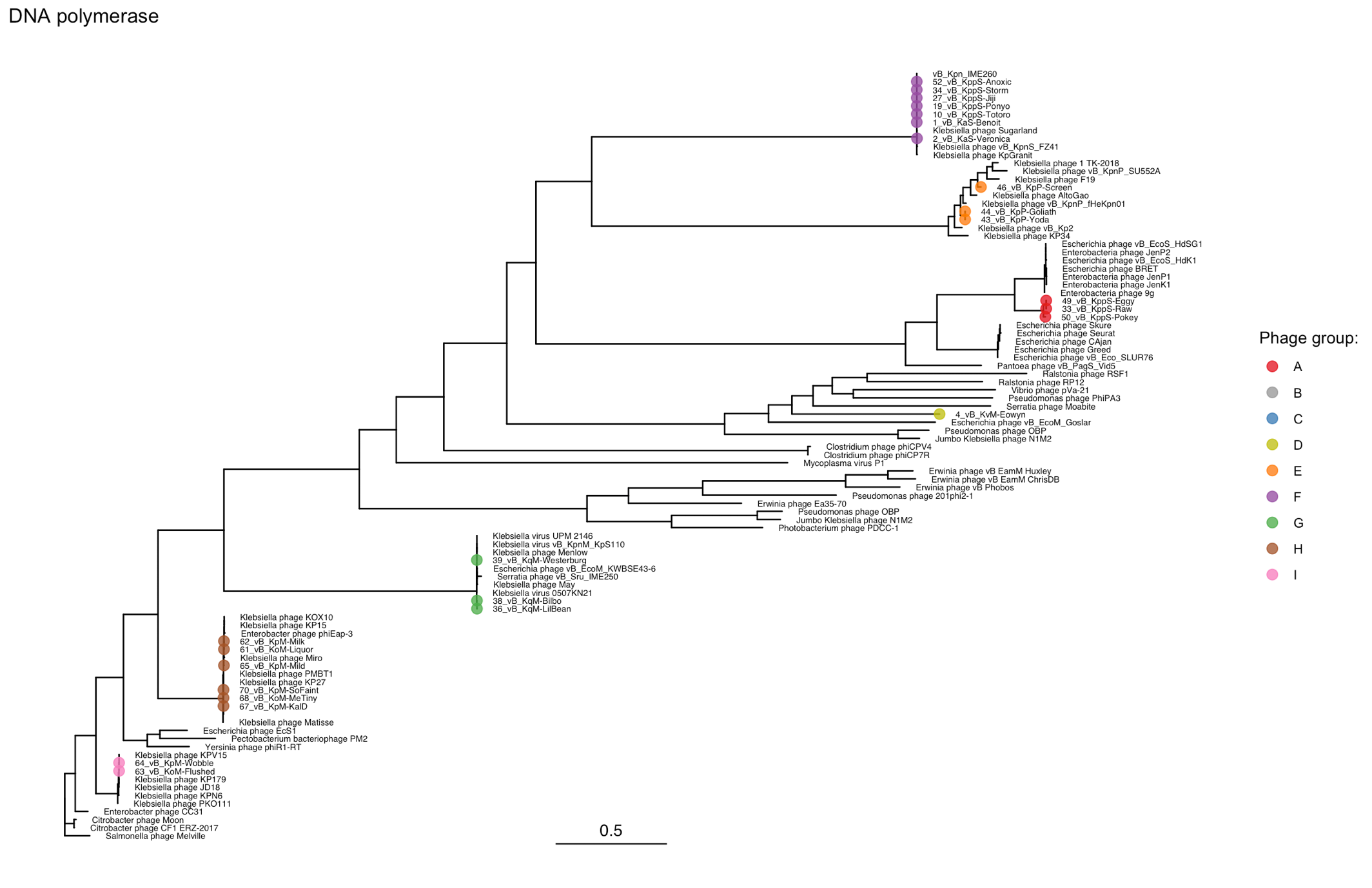

### Supplementarty fig S12

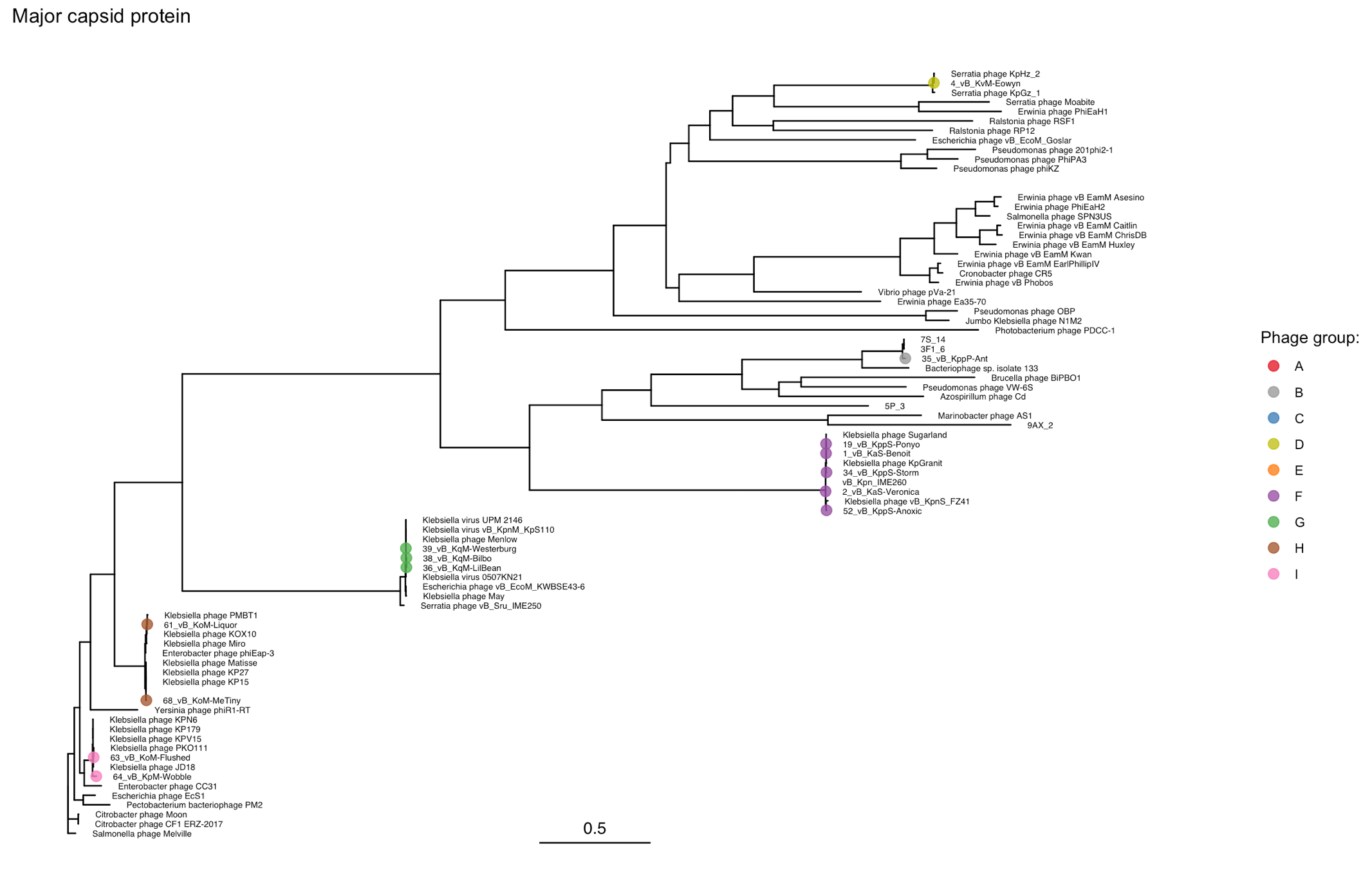

### Supplementarty fig S13

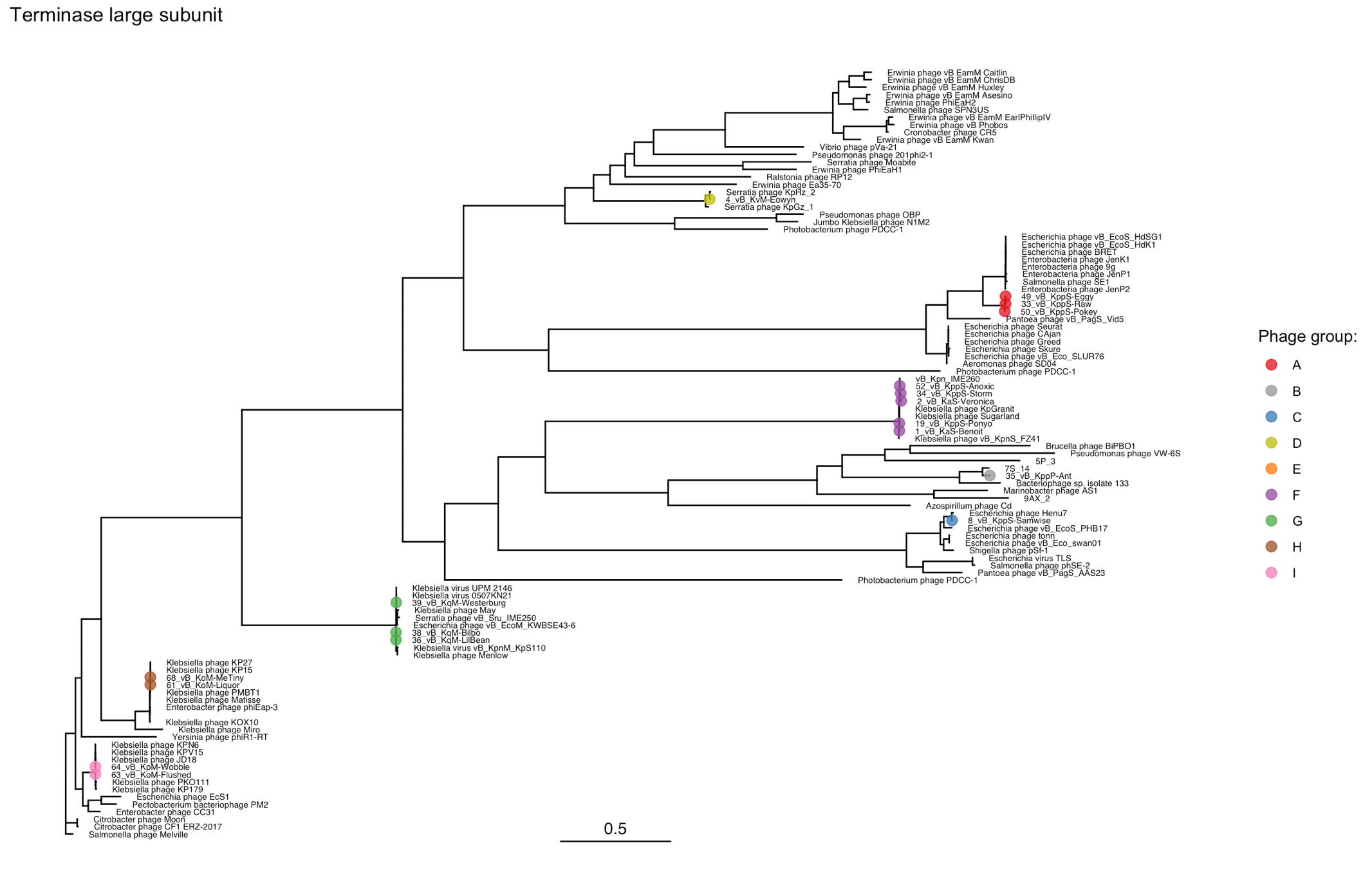
