## Supplementarty table S1 for "Isolation and characterisation of *Klebsiella* phages for phage therapy"

| Phage | Family | Genome Acc Number | Host Capsule Range | Capsule Depolymerase (Dp) | Dp Acc number | Dp Activity | Reference |
| --- | --- | --- | --- | --- | --- | --- | --- |
| 0507-KN2-1 | *Ackermannviridae* | AB797215 | KN2 | ORF96 | BAN78446 | KN2 | Hsu et al 2013 |
| NTUH-K2044-K1-1 | *Autographviridae* | AB716666 | K1 | KI-ORF34 | YP_009098385 | K1 | Lin et al 2014 |
| KP36 | *Drexlerviridae* | NC_029099 | K63 | depoKP36 (gp50) | YP_009226010.1 | K63 | Majkowska-Skrobek et al 2016 |
| K64-1 | *Myoviridae* | LC121097 | K1, K11, K21, K25, K30, K35, K64, K69, KN4, KN5 | S1-1  S1-2  S1-3  S2-1  S2-2  S2-3  S2-4  S2-5  S2-6 | BAW85694  BAW85692  BAW85693  BAW85695  BAW85696  BAW85697  BAW85698  BAQ02780  BAW85699 | K11  KN4  K21  KN5  K25  K35  K1  K64  K30, K69 | Pan et al 2017 |
| K5-2 | *Autographviridae* | KY389315 | K5, K24, K30, K38, K40, K52, K69 | K5-2 (ORF37)  K5-2 (ORF38) | APZ82804  APZ82805 | K30, K69  K5 | Hsieh et al 2017 |
| K5-4 | *Autographviridae* | KY389316 | K5, K8 | K5-4 (ORF37)  K5-4 (ORF38) | APZ82847  APZ82848 | K8  K5 | Hsieh et al 2017 |
| KpV71 | *Autographviridae* | KU666550 | K1, K62 | kpv71_52 | AMQ66478 | K1 | Solovieva et al 2018 |
| KpV74 | *Autographviridae* | KY385423 | K2, K13 | kpv74_56 | APZ82768.1 | K2, K13 | Solovieva et al 2018 |
| KP32 | *Autographviridae* | NC_013647 | K3, K21 | KP32gp37  KP32gp38 | YP_003347555  YP_003347556 | K3  K21 | Majkowska-Skrobek et al 2018 |
| vB_KpnP_IME321 | *Autographviridae* | MH587638 | KN1 | Dp42 | AXE28435 | KN1 | Wang et al 2019 |
| KN1-1 | *Autographviridae* | LC413193 | KN1 | KN1dep | BBF66844 | KN1 | Pan et al 2019 |
| KN3-1 | *Autographviridae* | LC413194 | KN3, K56 | KN3dep  K56dep | BBF66867  BBF66868 | KN3  K56 | Pan et al 2019 |
| KN4-1 | *Autographviridae* | LC413195 | KN4 | KN4dep | BBF66888 | KN4 | Pan et al 2019 |
| SH-KP152226 | *Autographvirinae* | MK903728 | K47 | Dep42 | QDF14644 | K47 | Wu et al 2019 |
| IME205 | *Autographvirinae* | KU183006 | K47 | Dpo42  Dpo43 | ALT58497  ALT58498 | K47  K47 | Liu et al 2020 |

Table SX – List of characterised *Klebsiella* targeting depolymerase tail-fibre proteins used to construct the phylogenetic tree (Figure SX).
