## Supplementarty table S2 for "Isolation and characterisation of *Klebsiella* phages for phage therapy"

| Phage | Family | Genus | Putative depolymerase | Protein Length (aa) | BLAST, HMMer, HHpred descriptions | HHpred Predicted boundaries (aa) | HHpred score (E-value) | PDB code |
| --- | --- | --- | --- | --- | --- | --- | --- | --- |
| vB_KvM-Eowyn | *Myoviridae* | *Iapetusvirus* | gp225 | 1183 | Bacteriophage CBA120 tailspike-protein4; hydrolase; Tail_spike_N  Tailspike domain protein gp42, Peptidase_S74, chaperone | 30-502  962-1117 | 0.0017  1.6e-11 | 5W6H  6EU4 |
|  |  |  | gp227 | 591 | Tail fiber protein, β-helical, pectate lyase  Particle associated glycoside hydrolase | 3-428  3-431 | 8.2e-21  2.0e-20 | 5W5P  6C72 |
|  |  |  | gp230 | 581 | K5 lyase  Tailspike-protein, beta-helix | 10-318  42-560 | 7.5e-14  7.1e-13 | 2X3H  4XOT |
|  |  |  | gp233 | 820 | Putative endo-N-neuraminidase; Outer surface protein A, hydrolase  K5 lyase | 1-420  57-548 | 5.6e-17  5.5e-15 | 5ZRU  2X3H |
|  |  |  | gp235 | 556 | Bacteriophage CBA120 tailspike-protein4; hydrolase  Tailspike-protein; parallel beta helix, putative endo-glycosidase | 12-533  12-400 | 1.2e-19  1.4e-15 | 5W6H  6NW9 |
|  |  |  | gp237 | 616 | Endo-xylogalacturonan hydrolase; Putative endo-N-acetylneuraminidase | 275-500 | 26 | 4CL2 |
|  |  |  | gp239 | 793 | Plasmin and fibronectin-binding protein A  Beta_Helix; Putative tail fiber; tailspike; hydrolase CBA120 | 83-582  40-521 | 7.8e-24  5.7e-21 | 4MR0  5W6S |
| vB_KppS-Raw | *Siphoviridae* | *Nonavirus* | gp50 | 740 | Pectate_lyase_3 superfamily; Beta_helix, phage_tail_N;  Outer Surface Protein A; hydrolase  Plasmin and fibronectin-binding protein | 208-730  291-732 | 5.4e-21  6.1e-19 | 5ZRU  4MR0 |
| vB_KppS-Eggy | *Siphoviridae* | *Nonavirus* | gp67 | 740 | Pectate_lyase_3 superfamily; Beta_helix, phage_tail_N;  Outer Surface Protein A; hydrolase  Plasmin and fibronectin-binding protein | 208-730  291-732 | 5.4e-21  6.1e-19 | 5ZRU  4MR0 |
| vB_KppS-Pokey | *Siphoviridae* | *Nonavirus* | gp78 | 740 | Pectate_lyase_3 superfamily; Beta_helix, phage_tail_N;  Outer Surface Protein A; hydrolase  Plasmin and fibronectin-binding protein | 205-730  291-732 | 1.1e-21  5.5e-19 | 5ZRU  4MR0 |
| vB_KpvM-LilBean | *Ackermannviridae* | *Taipeivirus* | gp52 | 1039 | phiAB6 tailspike; beta helix; superhelical trimer  Tailspike protein; parallel beta helix, putative endo-glycosidase | 86-813  209-851 | 1.0e-21  6.4e-17 | 5JS4  6NW9 |
|  |  |  | gp54 | 728 | phiAB6 tailspike; beta helix; superhelical trimer  Tailspike protein; parallel beta helix, putative endo-glycosidase | 1-552  140-576 | 1.5e-19  1.2e-17 | 5JS4  6NW9 |
|  |  |  | gp56 | 735 | Tailspike protein  Beta-1,3-glucanase; cellulose, glucanase  Beta-1,3-glucanase; tandem beta-helix; glucosidase, hydrolase | 81-282  75-282 | 4.0e-16  9.0e-16 | 5M5Z  3EQN |
|  |  |  | gp58 | 663 | Tailspike protein; parallel beta helix, putative endo-glycosidase  Bacteriophage CBA120 tailspike-protein4; hydrolase | 1-490  18-480 | 8.2e-13  1.2e-11 | 6NW9  5W6H |
| vB_KpvM-Bilbo | *Ackermannviridae* | *Taipeivirus* | gp55 | 1039 | phiAB6 tailspike; beta helix; superhelical trimer  Tailspike protein; parallel beta helix, putative endo-glycosidase | 86-813  209-851 | 1.0e-21  6.4e-17 | 5JS4  6NW9 |
|  |  |  | gp57 | 728 | phiAB6 tailspike; beta helix; superhelical trimer  Tailspike protein; parallel beta helix, putative endo-glycosidase | 1-552  140-576 | 1.5e-19  1.2e-17 | 5JS4  6NW9 |
|  |  |  | gp59 | 747 | Particle-associated glycoside hydrolase; glycosidase; tailspike  Tail fiber protein; AM27, tailspike protein | 12-354  11-320 | 6.9e-14  1.7e-12 | 6C72  5W5P |
|  |  |  | gp61 | 663 | Tailspike protein; parallel beta helix, putative endo-glycosidase  Bacteriophage CBA120 tailspike-protein4; hydrolase | 1-490  18-480 | 8.2e-13  1.2e-11 | 6NW9  5W6H |
| vB_KpvM-Westerburg | *Ackermannviridae* | *Taipeivirus* | gp161 | 1039 | phiAB6 tailspike; beta helix; superhelical trimer  Tailspike protein; parallel beta helix, putative endo-glycosidase | 86-813  371-852 | 3.7e-20  1.8e-16 | 5JS4  6NW9 |
|  |  |  | gp163 | 735 | Bacteriophage CBA120 tailspike-protein4; hydrolase; Tail_spike_N  phiAB6 tailspike; beta helix; superhelical trimer  CBM22; Binding site, carbohydrates, enzyme stability | 77-735  1-492  577-735 | 1.7e-31  7.1e-15  0.00017 | 5W6H  5JS4  4XUO |
|  |  |  | gp165 | 284 | Tailspike protein; parallel beta helix, putative endo-glycosidase  Rhamngalacturonase A; hydrolase; parallel beta-helix glycosidase | 1-277  98-180 | 2.3e-9  5.5e-8 | 6NW9  1RMG |
|  |  |  | gp166 | 552 | Polygalacturonase, glycosylhydrolase  Endopolygalacturonase; beta helical structure; glycoside hydrolase | 6-280  7-280 | 3.1e-10  3.3e-10 | 1IB4  1K5C |
|  |  |  | gp168 | 675 | Depolymerase KP32gp38; Klebsiella pneumoniae capsule depolymerase  Tailspike protein; parallel beta helix, putative endo-glycosidase | 93-675  1-418 | 5.6e-107  5.6e-19 | 6TKU  6NW9 |
| vB_KpP-Yoda | *Autographiviridae* | *Druliivirus* | gp52 | 464 | Bacteriophage CBA120 tailspike-protein4; hydrolase  Tailspike protein gp42  Endo-1,4-Beta-Xylanase Y; carbohydrate binding module | 4-464  2-267  307-464 | 1.0e-28  4.5e-11  0.0026 | 5W6H  6EU4  1DYO |
| vB_KpP-Goliath | *Autographiviridae* | *Druliivirus* | gp54 | 577 | Bacteriophage CBA120 tailspike-protein4; hydrolase  Tailspike protein gp42  Endo-1,4-Beta-Xylanase Y; carbohydrate binding module | 1-576  8-407  420-576 | 1.2e-35  8.3e-25  0.0068 | 5W6H  6EU4  1DYO |
| vB_KpP-Screen | *Autographiviridae* | *Druliivirus* | gp01 | 577 | Bacteriophage CBA120 tailspike-protein4; hydrolase  Tail fiber protein; AM27 Tailspike protein  Endo-1,4-Beta-Xylanase Y; carbohydrate binding module  Phage_Tail_middle | 3-576  8-355  420-577 | 1.1e-37  3.6e-24  0.003 | 5W6H  5W5P  1DYO |
