## Supplementarty table S3 for "Isolation and characterisation of *Klebsiella* phages for phage therapy"

| Phage | Lab ID | Tail length (nm) | Capsid width (nm) | Morphology | Genome size bp |
| --- | --- | --- | --- | --- | --- |
| vB_KppS-Raw | 33 | 153 | 46 | siphovirus | 61,195 |
| vB_KppS-Eggy | 49 | 141 | 55 | siphovirus | 60,681 |
| vB_KppS-Pokey | 50 |  |  | siphovirus | 61,218 |
| vB_KppS-Ant | 35 |  |  |  | 16,548 |
| vB_KaS-Gatomon | 6 | 60 | 54 | siphovirus | 49,702 |
| vB_KaS-Ahsoka | 7 | 200 | 60 | siphovirus | 49,702 |
| vB_KppS-Samwise | 8 | 204 | 65 | siphovirus | 49,891 |
| vB_KvM-Eowyn | 4 | 171 | 130 | myovirus | 268,550 |
| vB_KpP-Yoda | 43 | 13 | 51 | podovirus | 44,122 |
| vB_KpP-Goliath | 44 | 10 | 41 | podovirus | 44,010 |
| vB_KpP-Screen | 46 | 14 | 53 | podovirus | 43,749 |
| vB_KaS-Benoit | 1 | 180 | 55 | siphovirus | 109,014 |
| vB_KaS-Veronica | 2 | 160 | 60 | siphovirus | 110,196 |
| vB_KppS-Totoro | 10 | 170 | 70 | siphovirus | 109,014 |
| vB_KppS-Ponyo | 19 | 240 | 70 | siphovirus | 109,014 |
| vB_KppS-Jiji | 27 | 220 | 73 | siphovirus | 113,155 |
| vB_KppS-Storm | 34 | 203 | 69 | siphovirus | 110,834 |
| vB_KppS-Anoxic | 52 |  |  | siphovirus | 109,500 |
| vB_KqM-LilBean | 36 | 124 | 82 | myovirus | 158,859 |
| vB_KqM-Bilbo | 38 | 118 | 78 | myovirus | 158,858 |
| vB_KqM-Westerburg | 39 | 124 | 90 | myovirus | 158,883 |
| vB_KoM-Pickle | 12 | 121 | 65 | myovirus | 175,221 |
| vB_KoM-Liquor | 61 |  |  | myovirus | 176,734 |
| vB_KpM-Milk | 62 | 127 | 70 | myovirus | 176,734 |
| vB_KpM-Mild | 65 | 107 | 82 | myovirus | 176,856 |
| vB_KpM-KalD | 67 | 124 | 93 | myovirus | 174,351 |
| vB_KoM-MeTiny | 68 | 99 | 67 | myovirus | 175,419 |
| vB_KpM-SoFaint | 70 | 120 | 78 | myovirus | 175,933 |
| vB_KoM-Flushed | 63 | 115 | 70 | myovirus | 168,773 |
| vB_KpM-Wobble | 64 | 83 | 83 | myovirus | 168,530 |
